# Regulatory effects of the *Uty*/*Ddx3y* locus on neighboring chromosome Y genes and autosomal mRNA transcripts in adult mouse non-reproductive cells

**DOI:** 10.1101/2020.06.30.180232

**Authors:** Christian F. Deschepper

**Affiliations:** Cardiovascular Biology Research Unit, Institut de recherches cliniques de Montréal (IRCM) and Université de Montréal

## Abstract

In addition to sperm-related genes, the male-specific chromosome Y (chrY) contains a class of ubiquitously expressed and evolutionary conserved dosage-sensitive regulator genes that include the neighboring *Uty, Ddx3y* and (in mice) *Eif2s3y* genes. However, no study to date has investigated the functional impact of targeted mutations of any of these genes within adult non-reproductive somatic cells. We thus compared adult male mice carrying a gene trap within their *Uty* gene (*Uty^GT^*) to their wild-type (WT) isogenic controls, and performed deep sequencing of RNA and genome-wide profiling of chromatin features in extracts from either cardiac tissue, cardiomyocyte-specific nuclei or purified cardiomyocytes. The apparent impact of *Uty^GT^* on gene transcription concentrated mostly on chrY genes surrounding the locus of insertion, *i.e. Uty, Ddx3y*, long non-coding RNAs (lncRNAs) contained within their introns and *Eif2s3y*, in addition to possible effects on the autosomal *Malat1* lncRNA. Notwithstanding, *Uty^GT^* also caused coordinate changes in the abundance of hundreds of mRNA transcripts related to coherent cell functions, including RNA processing and translation. The results altogether indicated that tightly co-regulated chrY genes had nonetheless more widespread effects on the autosomal transcriptome in adult somatic cells, most likely due to mechanisms other than just transcriptional regulation of corresponding protein-coding genes.

## INTRODUCTION

Although the contributions of genes on the male sex chrY have long believed to be restricted to their effects on reproductive functions in sex organs, increasing evidence indicates that their impact may in fact extend to somatic cells as well. Thus, several types of genetic rodent models have shown that chrY and/or some of its genetic variants affect a range of phenotypes within the cardiovascular, immune, metabolic or neurologic systems^1,2,3,4,5^. In humans, one specific haplotype of the male-specific portion of chrY (MSY) associates with increased coronary artery disease risk, along with altered gene expression within macrophages^6^, while mosaic loss of chrY in blood cells (a relatively frequent and age-related event) associates with a host of disease manifestations, including shorter survival, increased cancer risk, cardiovascular events, Alzheimer disease and age-related macular degeneration^7,8,9,10^.

In addition of the male sex-determination *Sry* gene, MSY carries only a handful of protein-coding genes, which fall for the most part in two classes. The first one corresponds to genes expressed only in male reproductive tissues and/or specializing in sperm-related functions^11,12^. Genes within the second class share a particular set of properties, as they: *i*) are expressed ubiquitously in all tissues; *ii*) are X-degenerate genes that have on chromosome X (chrX) counterparts that escape chrX-inactivation in females; *iii*) exert their effects in a dose-related fashion that depends on expression of copies from both sex chromosomes; and *iv*) belong to an evolutionary conserved core of genes found in MSY from all placental vertebrates^12^. Although there are some species-specific variation concerning the exact nature of genes found in that shared core group, some of them (including *Uty, Ddx3y* and *Kdm5d*) are common to all investigated species^12^.

In addition to the above properties, the X-degenerate genes on MSY have functional annotations that relate mostly to some of the most basic cell functions (such as chromatin modification, ubiquitination, translation, splicing and chromatin modification). To date, there is evidence (albeit limited) that X-degenerate genes on MSY show at least some level functional redundancy with their homologs on chrX. For instance, *DDX3Y* has been shown to be functionally interchangeable with *DDX3X* in cultured mammalian cells^13,14^. Likewise, studies using mice with inactivating gene traps of either *Utx* or *Uty* demonstrated that *Uty* showed high level of functional redundancy with *Utx* during development, with further tests showing that the effects of both genes were independent of any lysine demethylase activity^15^. In humans, overexpressed *UTY* has been shown to rescue preleukemic phenotypes induced by *UTX* deficiency, also in a manner that was independent of any lysine demethylase activity^16^. However, there is still no study to date exploring whether targeted mutations of MSY genes do have an impact on somatic non-reproductive cells in adult organisms. To address this particular question, we used male mice carrying an inactivating gene trap within the fourth intron of *Uty* (*Uty^GT^* in short). These mice were previously shown to be fully viable^15^, but not used for studying phenotypes other than those occurring during the developmental stage. In particular, we performed deep sequencing of RNA (RNA-Seq) in hearts and cardiomyocytes from either adult *Y^UtyGT^* mice or their isogenic wild-type (WT) controls. The results were compared to those obtained with either a chromatin immunoprecipitation and sequencing (ChIP-Seq) assay of Histone H3 acetyl lysine-27 (H3K27ac) residues (*i.e*. marks of active enhancers) or with assays for transposase-accessible chromatin using sequencing (ATAC-Seq), which quantifies chromatin accessibility across the genome. We found that despite significant effects on hundreds of mRNA transcripts in cardiac cells, the genomic regulatory effects *Uty^GT^* were much more restricted, as they concentrated mostly on genomic regions on MSY. The data suggested that while the transcriptional effects of the *Uty* locus were restricted to a handful of regulators, the latter could nonetheless further regulate the transcriptomic profile of cells, presumably *via* post-transcriptional or post-translational mechanisms.

## RESULTS

### Regulatory effects of Uty^GT^ on surrounding MSY genes and genomic sequences

Combined analysis of the results of RNA-Seq, ChIP-Seq and ATAC-Seq demonstrated that the greatest impact of *Uty^GT^* was observed on either the abundance of transcripts from surrounding genes or chromatin marks in corresponding genomic regulatory regions. RNA-Seq revealed that *Ddx3y* and *Eif2s3y* were among the top most affected genes, either in terms of statistical significance or absolute fold change (Table 1). *Ddx3y* was the gene whose expression was (after *Uty* itself) affected most significantly, with its abundance in either hearts or cardiomyocytes from *Uty^GT^* being about ~ 15% of that in their WT counterparts (Table 1). *Eif2s3y* was among the 60 most affected genes but its expression was (contrary to *Ddx3y*) increased in hearts or cardiomyocytes from *Uty^GT^* mice. RT-qPCR experiments were performed to further validate those results. Since *Uty*, *Ddx3y* and *Eif2s3y* are all ubiquitously expressed MSY genes, non-cardiac tissues (including skeletal muscle, spleen, liver and/or primary lung fibroblasts) were included in the analysis. Regardless of the type of tissue, *Ddx3y* showed (similarly as detected by RNA-Seq) a greater than five-fold downregulation in samples from *Uty^GT^* mice (Fig. 1a). Likewise, *Eif2s3y* was upregulated in all tested tissues, the abundance of its mRNA transcripts in *Uty^GT^* tissues being about 150% of that found in their WT counterparts (Fig. 1b). In additional experiments, we verified whether *Uty^GT^* had any effect on expression of the chrX homologs of the above genes. Expression of *Utx* was unaffected and *Ddx3x* showed a slight increase of about 10 % (*P* < 0.05); expression of *Eif2s3x* was increased by about 40% (*P* < 0.01) (Supplementary Fig S1).

**Fig.1:**
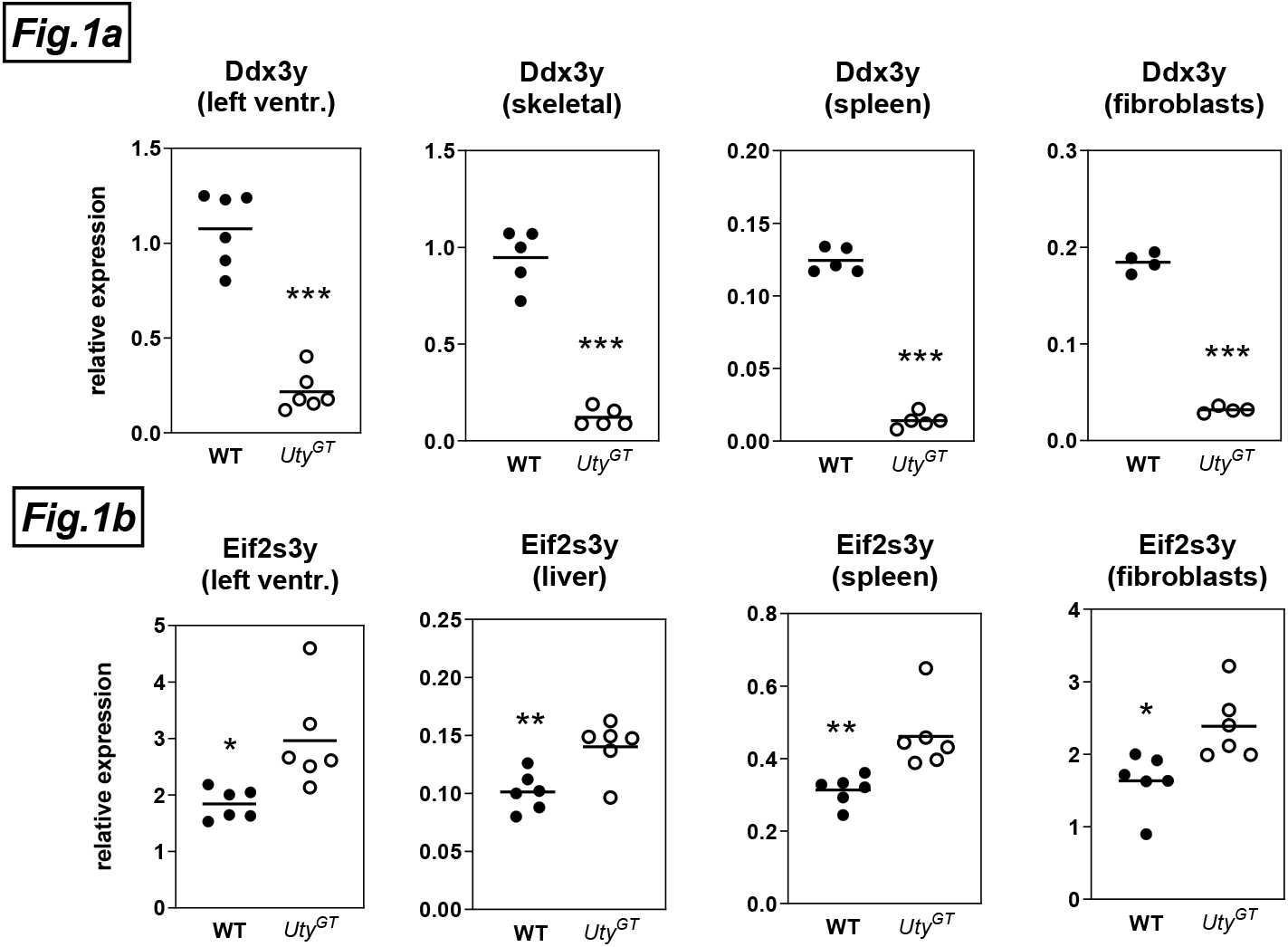
RT-qPCR quantification of abundance of mRNA transcripts of *Ddx3y* (Fig. 1a) or *Eif2s3y* (Fig. 1b). The experiments were performed using extracts from either cardiac left ventricles, skeletal muscle, liver, spleen or primary lung fibroblasts from either WT or *Uty^GT^* mice. Relative expression values corresponded to the ratios of Delta Ct values obtained for the gene of interest *vs*. those obtained for the *Rps16* normalizing gene.

**Table 1:** Effects of *Uty^GT^* on abundance of transcripts from MSY genes (KO/WT) and significance of the effects in either left ventricles or dispersed isolated adult cardiomyocytes.

|  | Left ventricles |  |  |  | Dispersed cardiocytes |  |  |
| --- | --- | --- | --- | --- | --- | --- | --- |
|  | *Uty* | *Ddx3y* | *Eif2s3y* |  | *Uty* | *Ddx3y* | *Eif2s3y* |
| **KO/WT** | 5.4% | 17.1% | 323% |  | 1.2% | 15.8% | 466% |
| **FDR** | 1.2 E-40 | 6.6 E-37 | 4.7E-06 |  | 1.0E-97 | 2.9E-89 | 9.7E-48 |
| **Rank FDR** | 1 | 2 | 59 |  | 1 | 2 | 4 |
| **Rank abs. FC** | 1 | 3 | 18 |  | 1 | 7 | 16 |

KO/WT: ratios of the FKPM values obtained for *Uty*<sup>GT</sup> mice vs. that of WT mice; FDR: false discovery rate; Rank FDR: relative rank for FDR values obtained for all tested genes; Rank abs. FC: relative rank for absolute fold values obtained for all tested genes

To further test whether the impact of *Uty^GT^* could be found at the level of proteins as well, we took advantage of the fact that the protein sequences of mouse Ddx3x and Ddx3y show 92% identity with each other as well as with human DDX3X or DDX3Y, and used two distinct antibodies against human recombinant DDX3X for Western blot analyses. Data obtained with each antibody were pooled by normalizing all data by the average of the values obtained in material from females (Fig. 2). Left ventricles from male XY^WT^ mice contained similar concentrations of overall Ddx3 immunoreactivity as that found in females; in contrast, the concentrations found in left ventricles from in male *XY^UtyGT^* mice was about 50% of that found in either male XY^WT^ or female XX mice. Similar differences were observed in primary fibroblasts from male XY^WT^ and *XY^UtyGT^* mice (Supplementary Fig S2). Altogether, the data indicated that the *Ddx3y* transcript was translated into a protein, and that its contribution to overall *Ddx3* immunoreactivity was about equal as that of its chrX homolog *Ddx3x*.

**Fig.2:**
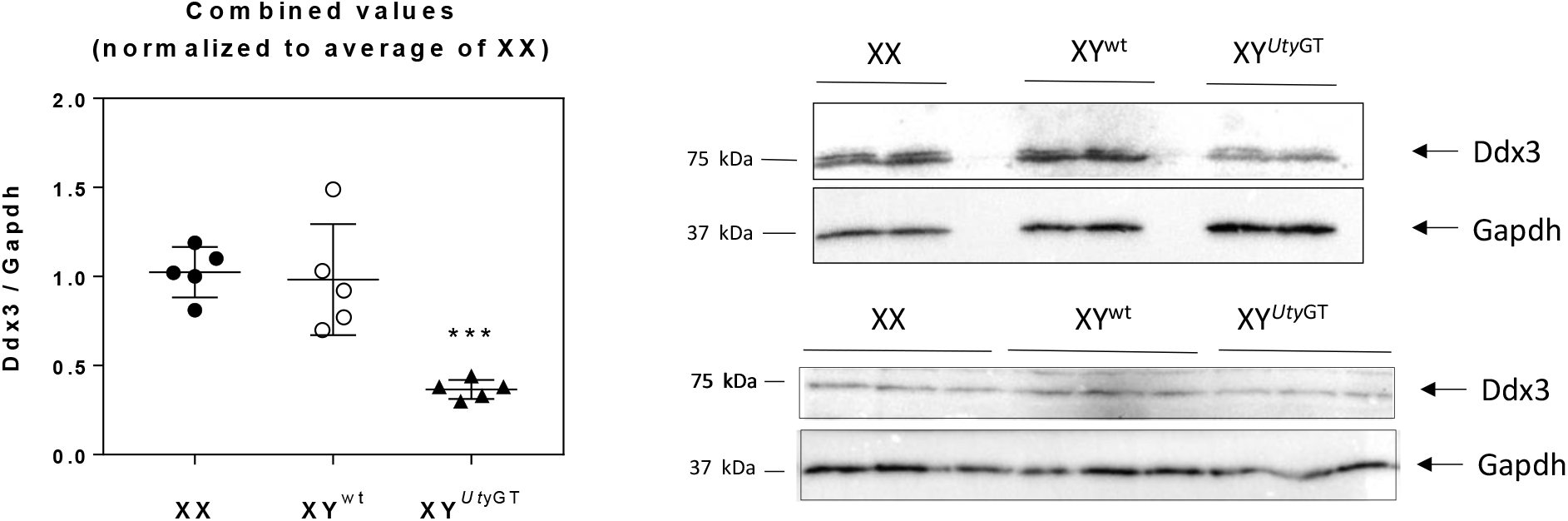
Western blot analyses of global Ddx3 immunoreactivity. The experiments were performed using left ventricular extracts from either female XX, male *XY^UtyWT^* or male *XY^UtyGT^* hearts. The experiments were performed using anti-DDX3 antibodies from either Santa Crux Biotechnologies (top right part) or Genetex (bottom right part. Results from each analysis were quantified and normalized by the average of values obtained in females and combined (right part). Full images of the hybridized membranes are provided in Supplementary Fig. S5 and Supplementary Fig. S6.

Genome-wide profiling of either H3K27ac marks or chromatin accessible regions was performed by ChIP-Seq and ATAC-Seq, respectively. In both cases, the most striking effects of *Uty^GT^* were observed in the vicinity of *Uty* and *Ddx3y*. With the H3K27ac ChIP-Seq, the regions showing the most highly significant differences in left ventricular chromatin from WT and *Uty^GT^* mice concerned two MSY peaks, *i.e*. one in the first exon region of *Uty* (FDR = 1.3E-27) and one in the first intron region of *Ddx3y* (FDR = 1.2E-27) (Fig.3). These peaks (detected in chromatin from WT mice, but not in that from their *Uty^GT^* counterparts) aligned with ENCODE-published adult mouse heart tracks for either H3K27Ac or p300 (two marks of active enhancers), DNase hypersensitive regions (another mark of accessible chromatin regions) or for Pol2 (a marker of active transcription initiation sites) (Fig. 3). The ATAC-Seq assays also detected differentially accessible regions in the proximal promoter region of *Uty* (FDR = 3.5 E-34) and the first intron of *Ddx3y* (FDR = 2.8 E-18) (i.e. regions overlapping with the positions of the two H3K27ac peaks described above), as well as in the proximal promoter of *Eif2s3y* (FDR = 5.50E-14) (Fig. 3).

**Fig.3:**
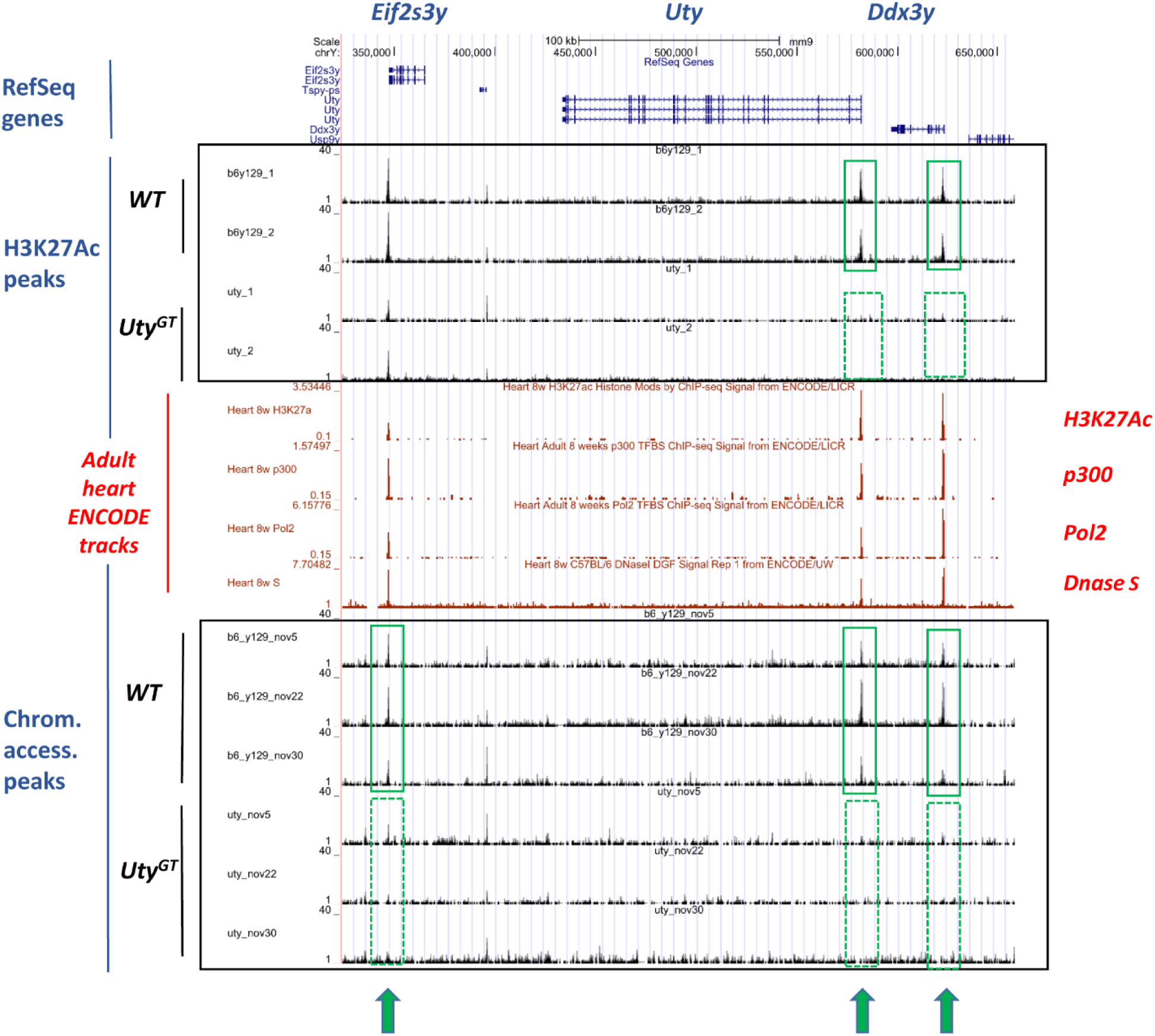
Graphic representation of results from the H3K27ac ChIP-Seq and ATAC-Seq analyses in MSY regions. Results represent those obtained within the genomic regions of the *Uty* locus and its two flanking genes. RefSeq gene coordinates are shown on top. Peaks for either H3K27ac ChIP-Seq or ATAC-Seq are shown in either the top or bottom framed areas, respectively. The areas framed in green indicate highlight peaks detected in WT, but not in *Uty^GT^* mice. All detected peaks align with those published by ENCODE (in red) and obtained using extracts from adult mouse hearts for either H3K27ac, p300 or Pol2 ChIP-Seq or Dnase hypersensitivity assays.

### Regulatory relationships between MSY genes

In primary lung fibroblasts from *Uty^GT^* mice, *Ddx3y* is down-regulated to the same extent as in heart or other non-cardiac tissues (Fig. 1a). To test whether such low *Ddx3y* levels could be rescued by re-initiation of expression of the protein-coding part of *Uty*, we thus infected such cells with viral particles carrying a lentiviral expression vector containing CMV-driven sequences for both *Uty* and the reporter eGFP gene, and isolated *Uty*-overexpressing cells by eGFP cytofluorometry. We found that *Uty* was clearly overexpressed in eGFP-positive cells, but that is was not associated with any changes in the low levels of Ddx3y expression (Fig. 4).

**Fig.4:**
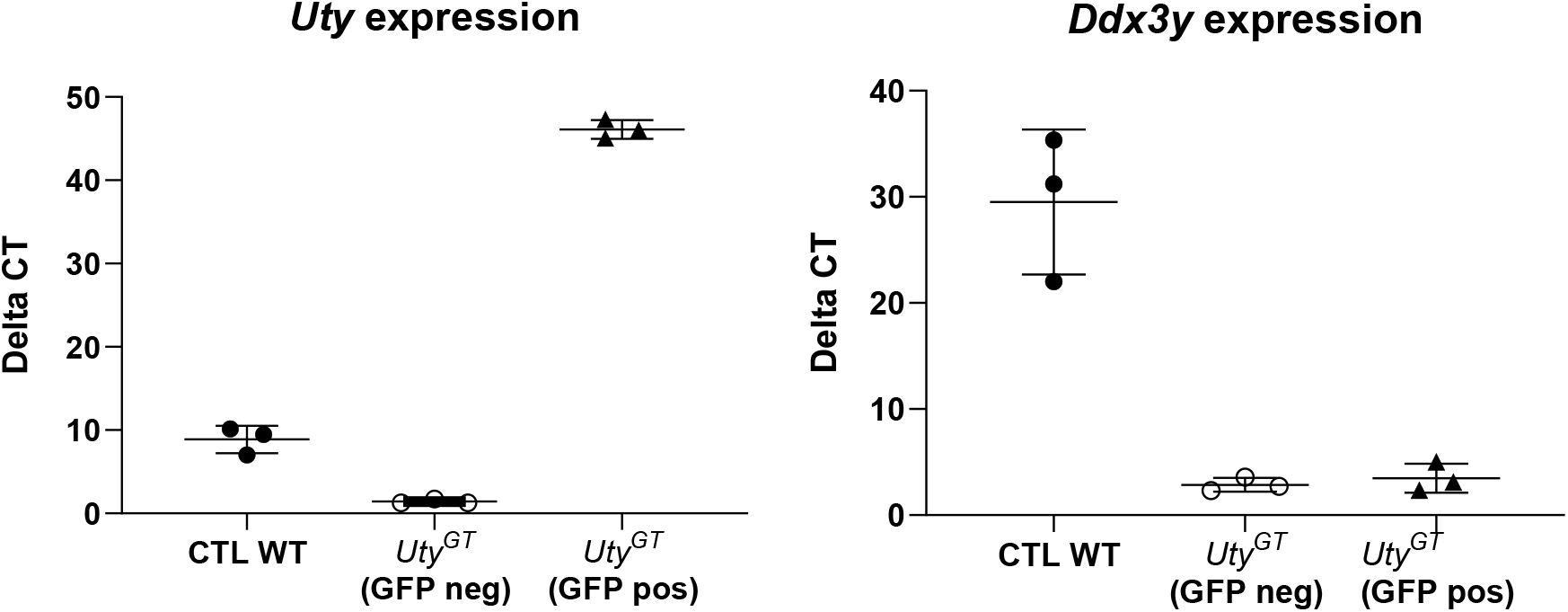
RT-qPCR quantification of abundance of mRNA transcripts of *Uty* (left) or *Ddx3y* (right). Experiments were performed in primary lung fibroblasts from *Uty^GT^* mice,em and infected with lentiviral particles containing lentiviral expression vectors that were either empty (CTL) or contained the full-length sequence of mouse *Uty*. In the latter case, the cells were sorted out by cytofluorometry to separate those with or without lentiviral integration (GFP positive or negative, respectively).

To test whether MSY genes could display high levels of co-expression constitutively, we used tools provided by the COEXPRESdb database to analyze data obtained on multiple tissues with a total of total 2236 Affymetrix arrays across 154 experiments^23^. Among all 21036 tested gene transcripts tested, *Uty* was found to interact most strongly with the protein coding *Ddx3y, Eif2s3y* and *Uba1y* genes, as well as with two other gene products: *i*) *Gm39552*, a three-exon lncRNA gene located within intron 25 of Uty; and *ii*) and *C030026M15Rik*, which corresponds to expressed sequences from Uty intron 14 (Supplemental Table 1). No autosomal gene showed co-expression scores approaching those found for MSY genes.

Since steady-state mRNA levels in cell cytoplasm are influenced by a variety of post-transcriptional mechanisms, it has been proposed that deep-sequencing of nuclear RNA (nucRNA-Seq) provides data that are more closely related to true transcriptional output^24,25^. We thus performed nucRNA-Seq on part of the nuclear material that was otherwise used for ATAC-Seq. Initial results indicated that the only transcripts affected significantly (FDR < 0.05) were those encoded by *Uty, Ddx3y*, and *Gm4017*, the latter being annotated as a processed pseudogene located within *Ddx3y* intron 17 (ENSMUSG00000101059). This intronic localization was reminiscent to that observed for *Gm39552* and *C030026M15Rik* (i.e. the two transcripts reported as co-expressed with *Uty* in the COEXPRESdb database). Visual inspection of the nucRNA-Seq sequencing tracks showed that these two transcripts could indeed be observed in samples from WT cardiomyocyte nuclei. However, unlike *Gm4017*, potential changes in their expression levels could not have been detected by the nucRNA-Seq as it was performed, because their coordinates were not included in the standard ENSEMBL mus GRCm38 gene build annotation file. We thus performed a novel nucRNA-Seq analysis that used a custom annotated file where the positions of *Gm39552* and *C030026M15Rik* were manually added to the GRCm38 gene build, in addition to that of *AK153586*, a previously detected expressed RNA transcript whose sequence matched genomic sequences within *Uty* intron 4 (Supplemental Fig. S3). When using these additional parameters, all transcripts emerged as significantly decreased (FDR < 0.05) in samples from *Uty^GT^* mice (Table 2). The presence of these transcripts in material from WT cardiomyocyte nuclei was further supported by the fact that FPKM values obtained in corresponding genomic regions were significantly higher (P < 0.05) than values found in remaining intron sequences from the genes they were inbedded in. In contrast, none of these transcripts were detected by the RNA-Seq analysis of cardiomyocyte extracts, where RNA originates predominantly from the cytoplasm.

**Table 2:** Top most differentially abundant RNA transcripts detected by the nucRNA-Seq analysis with custom annotation parameters.

| Ensembl gene ID | Gene name | mRNA | gene_biotype | localization | KO/WT | P-value | P-adj (FDR) |
| --- | --- | --- | --- | --- | --- | --- | --- |
| ENSMUSG00000068457 | Uty |  | protein_coding | MSY | 0.02 | 2.02E-135 | 6.03E-131 |
|  | AK153586 | AK153586 | expressed seq | Uty intron 4 | 0.00 | 1.08E-15 | 1.23E-11 |
|  | Gm39552 |  | lncRNA | Uty intron 25 | 0.00 | 1.24E-15 | 1.23E-11 |
|  | C030026M15Rik | AK049876 | lncRNA | Uty intron 14 | 0.00 | 1.54E-11 | 1.14E-07 |
| ENSMUSG00000069045 | Ddx3y |  | protein_coding | MSY | 0.16 | 1.97E-09 | 1.17E-05 |
| ENSMUSG00000101059 | Gm4017 |  | processed_pseudogene | Ddx3y intron 17 | 0.13 | 7.31E-05 | 3.63E-01 |
| ENSMUSG00000068745 | Mybphl |  | protein_coding | Chr3 | 0.26 | 1.65E-04 | 6.71E-01 |
| ENSMUSG00000069049 | Eif2s3y |  | protein_coding | MSY | 0.64 | 1.92E-04 | 6.71E-01 |
| ENSMUSG00000056673 | Kdm5d |  | protein_coding | MSY | 0.69 | 2.03E-04 | 6.71E-01 |

### Effects of Uty^GT^ on autosomal chromatin

Outside of MSY, the H3K27ac ChIP-Seq analysis detected only two autosomal regions with significant differences. The first one comprised three peaks on chr17 (with matching positions of ENCODE tracks for H3K27Ac, p300, Pol2 and DNase hypersensitivity) whose intensity was lower in *Uty^GT^* hearts than in their WT counterparts (Supplementary Fig. S4). However, neither RNA-Seq results nor additional RT-qPCR experiments performed on heart extracts (results not shown) showed any evidence of differential expression for any of the genes in the proximity of these peaks (*Dynlt1b, Tmem181a, Tmem181b*, and the lncRNA gene *Gm29719*). The second peak was found in a gene-poor intergenic region on chr15, with no matching peak within published ENCODE tracks.

With ATAC-Seq, differently accessible regions in autosomal chromatin were detected chr19 (in the intergenic regions between the long non-coding RNA (lncRNA) gene *Malat1* and *Neat1* (FDR = 9.6E-19) (Fig.5) (Supplemental Table 2). This signal was matched by increased expression of *Malat1* in *Uty^GT^* samples was found in heart samples by RNA-Seq, and further confirmed by RT-qPCR, both for heart as well as non-cardiac tissues (including skeletal muscle, liver and primary fibroblasts) (Fig. 6). We detected no change in expression of the *Malat1*-neighboring *Neat1* gene, either by RNA-Seq or additional RT-qPCR experiments (results not shown). Increased chromatin accessibility (FDR 1.E-31) was also found on chr5 in the region of the first intron of *En2*. However, we found no evidence indicating that the corresponding transcript was present in our samples, a finding in keeping with ENCODE transcriptome data showing that expression of this gene is mostly restricted to the developing central nervous system. In addition to *En2* and *Malat1*, some levels of differential accessibility were detected for about 100 other autosomal regions (Supplemental Table 2). However, the level of significance of other differentially accessibility regions dropped sharply compared to the top five most significantly affected regions, and there was no accompanying evidence for differential regulation of nearby genes.

**Fig.5:**
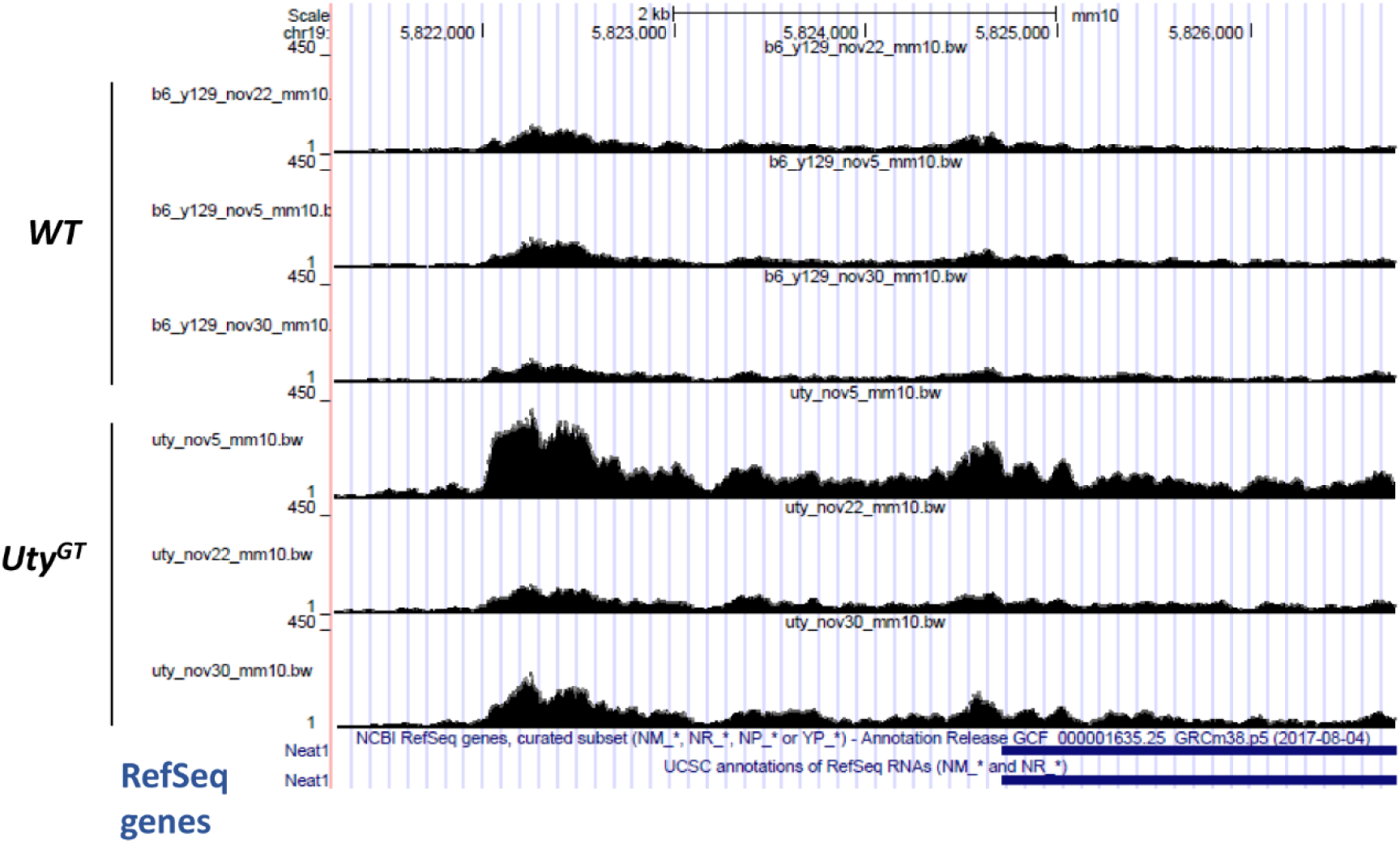
Graphic representation of results from ATAC-Seq analyses in chr19 regions. ATAC-Seq signals in cardiomyocyte nuclei from either WT or *Uty^GT^* mice (n = 3 in each group). The figure shows the intergenic region between the genes *Neat1* (partial coordinates shown at the bottom) and *Malat1* (located further downstream, coordinates outside of the boundaries of the window).

**Fig.6:**
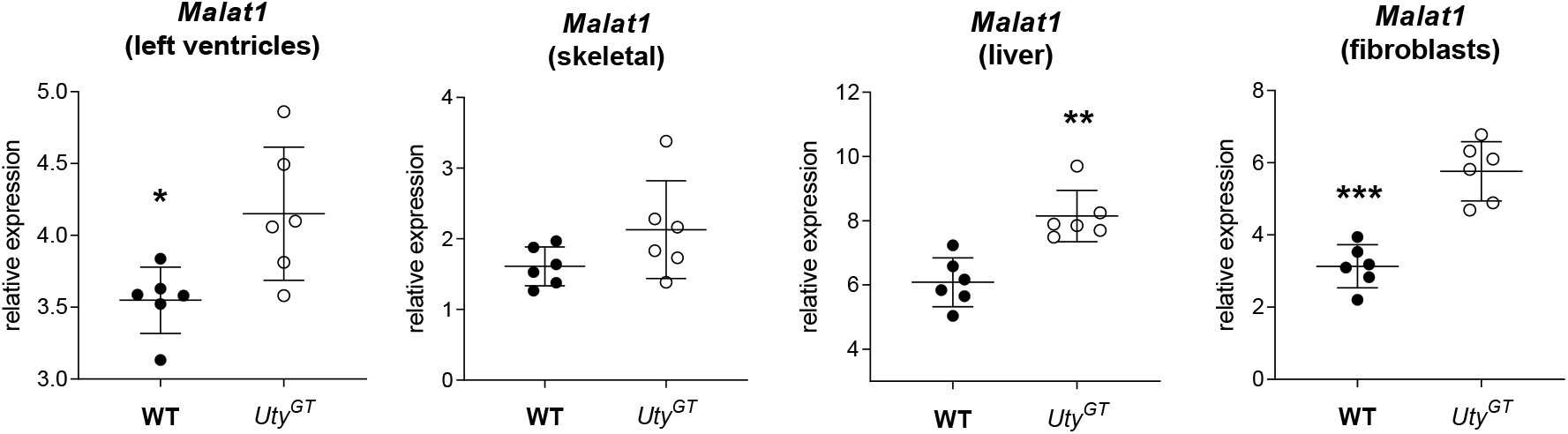
RT-qPCR quantification of abundance of *Malat1* mRNA transcripts in four different tissues from WT and *Uty^GT^* mice. *Malat1* expression is significantly increased (as determined by t-tests) in either left ventricles, liver and primary lung fibroblasts (*P < 0.05, **P < 0.01, ***P < 0.001, respectively) from *Uty^GT^* mice. A similar trend is observed in skeletal muscle, but did not reach statistical significance.

### Impact of Uty^GT^ on the adult heart transcriptome

The RNA-Seq experiments revealed that *Uty^GT^* was accompanied: *i*) in whole heart left ventricles, by down- and upregulation of 671 and 716 genes (FDR < 0.05), with absolute fold changes averaging 0.52 and 0.68 fold, respectively; and *ii*) in adult cardiocytes, by down- and upregulation of 669 and 476 genes (FDR < 0.05), with absolute fold changes averaging 0.49 and 0.63 fold, respectively. A total of 151 genes were common to both datasets. To test whether the transcriptomic changes could concern particular cellular functions, we looked for GO ontology terms showing significant enrichment (FDR < 0.01) among the *Uty^GT^*-affected genes (FDR < 0.05) with > 2-fold enrichment, then regrouped the terms meeting these criteria within broader categories that could encompass them. Among genes down-regulated in *Uty^GT^* mice, highly enriched GO terms could be consolidated with 2-3 overarching categories, accounting for 59% and 70% of GO terms enriched in differentially expressed genes in left ventricles and cardiomyocytes, respectively (Table 3)(Supplemental Table S6). Likewise, among genes up-regulated in *Uty^GT^* mice, enriched GO terms could be consolidated with 3-5 overarching categories, accounting for 45% and 50% of all enriched GO terms in left ventricles and cardiomyocytes, respectively (Table 4)(Supplemental Table S6). Both in ventricular extracts and isolated cardiomyocytes, genes up-regulated in *Uty^GT^* mice showed high and significant enrichment for ribosome-related GO terms.

**Table 3:**
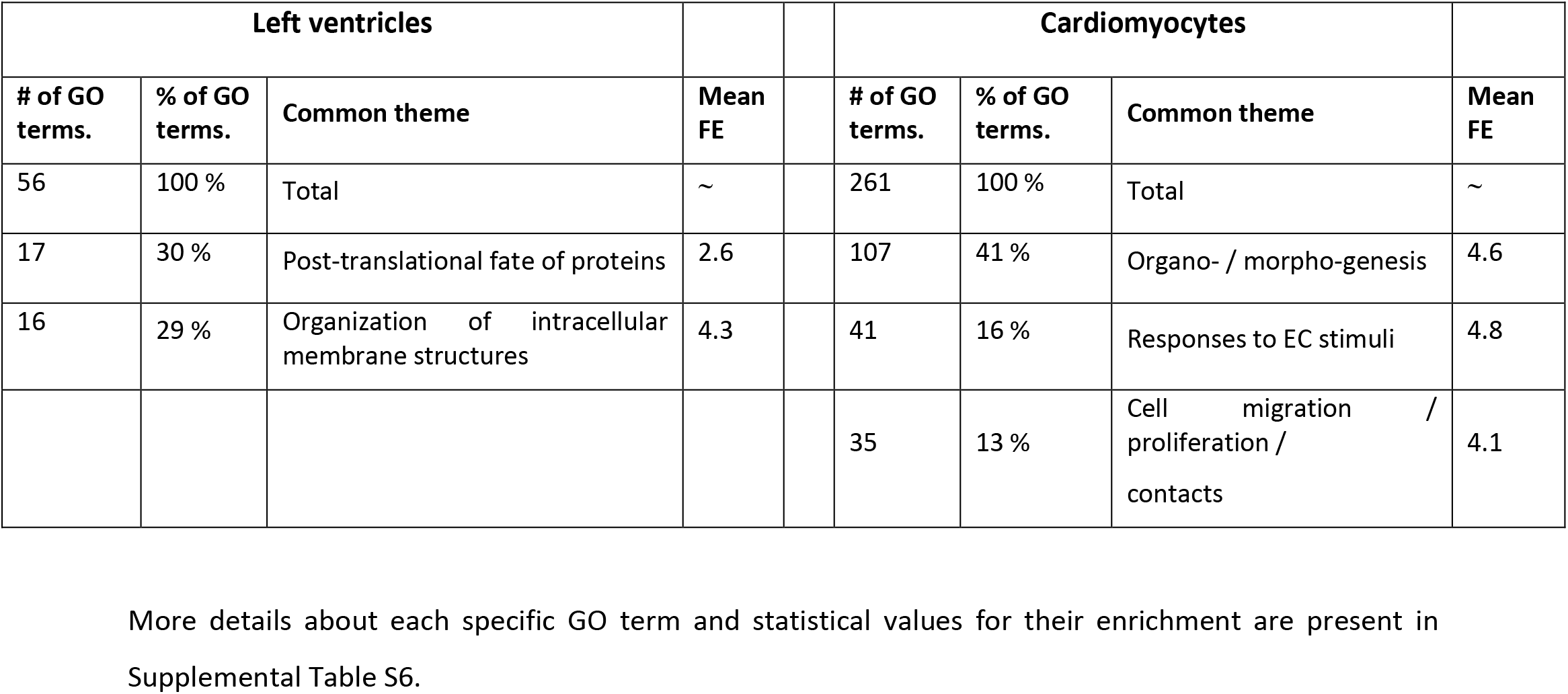
Common themes encompassing GO terms showing both high (> 2 fold) and significant (FDR < 0.01) enrichment among genes downregulated in *Uty^GT^* mice.

| Left ventricles |  |  |  |  | Cardiomyocytes |  |  |  |
| --- | --- | --- | --- | --- | --- | --- | --- | --- |
| # of GO terms. | % of GO terms. | Common theme | Mean FE |  | # of GO terms. | % of GO terms. | Common theme | Mean FE |
| 56 | 100 % | Total | ~ |  | 261 | 100 % | Total | ~ |
| 17 | 30 % | Post-translational fate of proteins | 2.6 |  | 107 | 41 % | Organo- / morpho-genesis | 4.6 |
| 16 | 29 % | Organization of intracellular membrane structures | 4.3 |  | 41 | 16 % | Responses to EC stimuli | 4.8 |
|  |  |  |  |  | 35 | 13 % | Cell migration / proliferation / contacts | 4.1 |

More details about each specific GO term and statistical values for their enrichment are present in Supplemental Table S6.

**Table 4:** Common themes encompassing GO terms showing both high (> 2 fold) and significant (FDR < 0.01) enrichment among genes upregulated in *Uty^GT^* mice.

| Left ventricles |  |  |  |  | Cardiomyocytes |  |  |  |
| --- | --- | --- | --- | --- | --- | --- | --- | --- |
| # of GO terms. | % of GO terms. | Common theme | Mean FE |  | # of GO terms. | % of GO terms. | Common theme | Mean FE |
| 111 | 100 % | Total | ~ |  | 224 | 100 % | Total | ~ |
| 20 | 18 % | Ribosomal activity | 6.7 |  | 42 | 18 % | Mitochondrial activity | 7.5 |
| 18 | 16 % | RNA processing | 3.3 |  | 36 | 16 % | Muscle contractile apparatus | 5.9 |
| 12 | 11 % | Chromosome maintenance | 3 |  | 14 | 6 % | Protein catabolism | 4.8 |
|  |  |  |  |  | 13 | 6 % | Ribosome organization | 7.3 |
|  |  |  |  |  | 13 | 6 % | Cytoskeleton |  |

More details about each specific GO term and statistical values for their enrichment are present in Supplemental Table S6.

## DISCUSSION

Despite mounting correlative evidence that MSY genes have effects outside of reproductive tissues, studies using gene targeting experiments to demonstrate in a direct fashion their impact within adult somatic cells and/or their mechanisms of action are still lacking. The current study demonstrates that a gene trap within the *Uty* gene does have demonstrable effects within adult somatic cell functions, but also that MSY genes may have some idiosyncratic properties. The most prominent effect of the *Uty^GT^* gene trap is that it is accompanied by a profound and ubiquitous downregulation of its neighboring *Ddx3y* gene and, by a smaller extent, by up-regulation of its other flanking gene (*Eif2s3y*). The dual effects of *Uty^GT^* on expression of both *Uty* and *Ddx3y* were accompanied by direct regulatory genomic changes involving a disappearance of both H3K27Ac and ATAC-Seq peaks in the proximal enhancer regions of these two genes. In contrast, the protein-coding product of the *Uty* gene did not appear to have direct regulatory actions on *Ddx3y* expression, because its re-expression in *Uty^GT^* primary cells did not alleviate the downregulation of *Ddx3y*. One possible explanation for these observations (for which there is actually some support within the literature) could be that non-coding sequences within the *Uty/Ddx3y* locus actually play a role in the co-regulation of these two genes. Indeed, comparisons of chrY sequences from 5 mammalian species have revealed that, despite highly divergent gene order among all of them, the *Uty/Ddx3y* gene pair is part of a microsyntenic region that has been preserved over 340 million years of independently sampled evolutionary history^26^. Other phylogenomic studies have shown previously that: *i*) such conserved ancient microsyntenic pairs arise when conservation of *cis*-regulatory sequences was required for co-regulation of the genes, and *ii*) such sequences were enriched within non-coding regions of the locus, and often resided within particularly large introns of the genes within the conserved pair^27^. In fact, our nucRNA-Seq analysis revealed that 4 transcripts corresponding to genomic sequences within 3 introns of *Uty* and one of *Ddx3y* could indeed be detected in nuclei from WT mice, with some of them having been previously been detected in transcriptomic studies performed across a wide range of tissues and whose results have been compiled in publically available databases. In the present study, we also observed that abundance of all these non-coding transcripts was greatly decreased in nuclei from *Uty^GT^* mice cardiomyocytes. It thus appeared that the disappearance of H3K27ac and ATAC-Seq peak in the proximal enhancer regions of *Uty* and *Ddx3y* was accompanied by downregulation of not only the protein-coding parts of these genes, but also of transcripts originating from sequences within their introns.

In a more general fashion, the co-expression databases showed that high levels of co-expression were a defining feature of MSY genes in general. As this occurred not only in mice, but in humans as well, it may constitute an evolutionary conserved characteristic of MSY genes in vertebrates. Of note, such patterns of co-expression between neighboring genes are rarely seen within vertebrate genomes, and do almost never occur at such high levels^28^. Nonetheless, such characteristics may have emerged within the male chrY because evolutionary forces that are different from those exerted on autosomes^29^ have resulted in different types of regulatory mechanism. One other evolutionary conserved feature appeared to be that, among the most highly co-expressed MSY genes, a high proportion of them [4 out of 8 of genes (50%) in mice, and 15 out of 23 (65%) in humans] were in fact non-coding. Interestingly, a recent study analyzing transcripts in brains from human and chimpanzee embryos discovered six novel non-annotated lncRNAs originating from chrY. Such RNA transcripts may thus constitute a class of agents that mediate some MSY effects or, alternatively, participate to the high levels of co-expression among MSY genes. Indeed, while three-dimensional looping events might be invoked in mice (where all X-degenerate MSY genes are contained within a relatively small 1 MB genomic region) as possible mechanisms underlying high levels of co-expression among MSY genes, such mechanisms are less likely to occur in humans, where the loci of MSY genes show much greater dispersion across much larger regions. Moreover, we found that the non-coding RNAs co-expressed with *Uty* were detected in samples from cardiomyocyte nuclei but not in those extracted from whole cardiomyocytes, indicating that these transcripts correspond in fact nuclear lncRNAs. The latter are known to participate (among other biological processes) to chromatin organization and accessibility, in part via recruitment of chromatin-modulating proteins to nearby chromatin regions^30^. In particular, given that *Uty^GT^* resides within the same *Uty* intron that harbors *AK153586*, decreased production of the latter could constitute one initial event triggering other chromatin rearrangements within the whole locus.

Although *DDX3Y* has been shown to be functionally interchangeable with *DDX3X* in cultured mammalian cells^13,14^, some controversy remains concerning the production of the cognate protein outside of reproductive tissues. Indeed, some have argued that the production of the DDX3Y protein was (in contrast to the ubiquitous expression of the gene) restricted to the male germline^31,32^. In contrast, other studies have detected DDX3Y peptides in either neural progenitor or hematopoietic stem cells^33,34,35^. Our Western blot analysis are in line with the latter studies, providing evidence that *Ddx3y* can indeed be translated into a protein in at least mouse somatic cells. Moreover, the downregulation of *Ddx3y* caused by *Uty^GT^* is accompanied by an equivalent reduction in overall Ddx3 immunoreactivity. The importance of this finding is many-fold. For instance, it has been shown that hematopoietic loss of DDX3X results is lethal in developing females, but viable in males^36^; however, without knowing whether production of the DDX3Y protein can occur in somatic cells, it was not possible to determine whether this compensatory effect of MSY could be due to DDX3Y itself. Likewise, since the actions of DDX3X are dosage-related, confirmation of the production of DDX3Y protein supports its possible contribution to dose-related effects. Of note, due to differences in their promoters, the *DDX3Y* and *DDX3X* genes have been shown to be independently regulated^13^, indicating that male organisms may regulate overall DDX3 immunoreactivity in part *via* mechanisms that are different from those in females.

When the results of H3K27ac ChIP-Seq, ATAC-Seq, RNA-Seq and nucRNA-Seq are all considered collectively, the genes contained within the *Uty/Ddx3y* locus were the only ones for which evidence for transcriptional regulation via direct regulatory genomic effects was entirely consistent across experiments. For *Malat1*, increased chromatin accessibility was matched by increased abundance of mRNA abundance in tissue extracts, but not in nuclear extracts. For *Eif2s3y*, the results were more difficult to interpret, as decreased chromatin accessibility in its proximal promoter region was matched by a trend towards decreased abundance in nuclear extracts (Table 2), which contrasted with a significant (FDR < 0.01) increase in the abundance of its transcript in tissues (see discussion below). Other indications of effects of *Uty^GT^* on either H3K27ac abundance or accessibility in autosomal chromatin were not matched by evidence for changes in the abundance of the transcripts of genes in corresponding regions. While the direct transcriptional effects of *Uty^GT^* thus appeared to be relatively restricted, the gene trap nonetheless had substantial effects on the transcriptome of cardiac cells, as hundreds of mRNA transcripts showed differential abundance. The magnitudes of the average fold-changes (ranging from 0.49 to 0.62) were relatively modest, but within the range of those reported by others that have studied sex biases in gene expression^37,38^. Moreover, there was functional coherence in the transcript abundance changes, as GO term analyses revealed that the majority of terms showing enrichment in differentially regulated genes showed great overlap within a few well-defined functional categories. For genes downregulated in left ventricles of *Uty^GT^* mice, these functions corresponded to post-translational protein processes (including protein localization, oligomerization and phosphorylation) and maintenance of intracellular membranes (including organelles, mitochondria and sarcolemma). For genes upregulated in left ventricles of *Uty^GT^* mice, these functions corresponded to ribosomal activity (including ribosomal organization and translation), processing and metabolism of RNA, and chromosome maintenance. Overall, these functional categories are compatible with the proposed concept that MSY X-degenerate genes are regulators of some of the most basic cellular homeostatic maintenance mechanisms^12^. More specifically, the categories related to ribosomal activity and RNA processing might partly explain the somewhat contradictory results obtained for *Eif2s3y*. Indeed, the RNA-Seq of left ventricular extracts indicated that, in addition to increased *Eif2s3y*, the transcripts of three other eukaryotic initiation factors (*Eif3a, Eif4e3*, and *Eif5*) were significantly increased, whereas those of three others (*Eif4g1, Eif3f*, and *Eif3h*) were significantly decreased (FDR < 0.01). Accordingly, *Uty^GT^* might affect the stability of several eukaryotic initiation factor transcripts, and affect that of *Eif2s3y* in a manner that is contrary to its effects on transcription of the gene.

Interestingly, at least two of the genes for which we observed evidence of increased transcription are known regulators of post-transcriptional mechanisms. On one hand, the DDX3 proteins are RNA helicases genes involved in all aspects of RNA metabolism, including mRNA nuclear export, translation and decay, and ribosome biogenesis^39, 40^. In particular, they affect protein translation by a variety of mechanisms that include interactions with translation initiation factors^41,42^, assembly of functional ribosomes^43^ and N^6^-methyladenosine modifications^44, 45^. On the other hand, *Malat1* may regulate: *i*) mRNA transcript stability via its ability to behave as a miRNA-binding competitive endogenous RNA; and *ii*) regulate the activity of proteins and/or assist in their cellular localization via its direct protein-binding capabilities^46^. All these functions are related to the GO categories showing highest levels of enrichment among *Uty^GT^*-regulated gene transcripts (*i.e*. to ribosomal activity and RNA processing). The above two genes may even work in complementary fashion, as DDX3 regulates the abundance of N^6^-methyladenosine residues in *Malat1*, which in turn may modulate its biological activity^44,47^.

Altogether, the current study shows that MSY genes have demonstrable effects on regulators of adult somatic cell functions. However, such effects might result from MSY behaving as an entire integrated regulatory unit rather than from individually regulated MSY genes, and might involve mechanisms other than just transcriptional regulation of autosomal protein-coding genes. These findings could constitute a stepping stone to better understand in which fashion chrY-related genetic differences may affect health-related outcomes.

## ACKNOWLEDGEMENTS

The work was partly funded by grant MOP-93583 from the Canadian Institutes for Health Research (CIHR). We are deeply grateful for the help of the IRCM Bioinformatics, Molecular Biology, Cytofluorometry and Animal Physiology Core Laboratories, and acknowledge in particular the contributions of Virginie Calderon, Caroline Grou, Odile Neyret, Myriam Rondeau, Éric Massicotte, Julie Lord and Manon Laprise. We also thank Sylvie Picard for technical help, and Mariana Bego and Mélanie Laporte for help with generation of lentiviral particles, and Pierre Bensidoun for immunocytochemistry and microscopy.

## MATERIAL AND METHODS

### Mouse strains and cells

All procedures involving mice were conducted following approval by the animal ethic committee of the Institut de Recherches Cliniques de Montréal (IRCM) and in agreement with the guidelines of the Canadian Council for Animal Care. *Uty*^*Gt*(*XS0378*)*Wtsi*^ (*Y*^*UtyGT*^ in short) ES cells (derived from the original parental E14Tg2a.4 ES cell line from 129P2/OlaHsd mice) were obtained from the Mutant Mouse Resource & Research Centers (MMRRC) and injected into C57BL/6J host blastocysts for chimera generation. Male chimeras were crossed to female C57BL/6J to assess germline transmission, and resulting males harboring the *Y^UtyGT^* were mated for more than 10 generations to female C57BL/6J mice. Isogenic control WT mice were generated by backcrossing 129P2/OlaHsd male mice to female C57BL/6J mice for the same number of generations. Primary fibroblasts were obtained from lungs of adult mice as described previously^49^. When cells reached confluence six days after initial plating, they were split 1:4 for passage 1 (P1) and allowed to reach confluence again. For experiments, all fibroblasts were passage 2. Mouse adult cardiocytes were obtained by digesting adult mouse hearts by retrograde perfusion of a solution of collagenase type 2 (Worthington, Lakewood, NJ) in calcium-free solutions, followed by gravity sedimentation within 5% bovine serum albumin (BSA) solutions supplemented in a step-wise fashion with increasing CaCl2 concentrations, as described ^50^. After isolation, the cells were plated on laminin-coated dishes, and used 4 hours after plating for RNA isolation.

### Isolation and flow-cytometry sorting of cardiomyocyte nuclei

Cardiomyocyte-specific nuclei were isolated from frozen left ventricular cardiac sample as described ^51,25^, with modifications to optimize maintenance of RNA integrity. In short, left ventricle tissue (priorly powdered under liquid nitrogen) was homogenized with a Polytron in a 40 mM Tris / 10 mM citrate buffer (to inhibit ribonucleases^52^) containing 5 mM KCl, 12% polyethylene glycol 8000 (to allow for isolation of nuclei in the absence of ribonuclease-activating ions ^53^), 1 mM DTT, 1 mM PMSF, protease inhibitors, 1.5 mM spermine and 20 mM sodium propionate (for chromatin integrity). The extracts were processed with a Dounce homogenizer and passed sequentially through 98 μm and 38 μm cell selectors. After spinning, the nuclei were resuspended in 1 ml of the same buffer supplemented with 3μl of an Alexa fluor 642-coupled antibody raised against the *pericentriolar 1* protein (PCM1) (SCBT, Dallas, TX), *i.e*. an antigen that accumulates specifically around nuclei from cardiomyocytes^51^. A sub-aliquot of that material was used for counterstaining with Hoechst bis-benzidine. After 30 min incubation at 4°C, the pellets were washed and resuspended in 1 ml of Tris/acetate buffered saline supplemented with 2% ribonuclease-free BSA, sorted out by cytofluorometry, and collected in tubes containing 200U of the Ribolock ribonuclease inhibitor (Thermofisher). The numbers of nuclei isolated in this manner were 50,000 and 150,000 for ATAC-Seq and RNA-Seq, respectively. Parallel analysis of the Hoechst-counterstained sub-aliquots showed that > 95% of PCM1 positive particles were Hoechst-positive nuclei, and comprised in average about 30% of all nuclei in the preparation.

### Sequencing-based procedures

Total RNA was isolated from tissues and cells using RNeasy micro kits with in-column DNase treatments (Qiagen, Toronto, Canada). For cardiac left ventricular samples (n = 7 for each mouse strain), preparation of 3’ end RNA sequencing libraries were prepared (using the Lexogen QuantSeq 3’ mRNA-Seq Library Prep Kit FWD library) and sequenced by the Neuroscience Genomics Core. Read-quality confirmation and alignment on the mouse GRCm38/mm10 genome were performed using Lexogen’s QuantSeq Bluebee analysis platform. For RNA from isolated adult cardiomyocytes (n = 3 for each mouse strain), ribodepletion was performed using the Fast Select mouse RNA removal kit (Qiagen), the RNA-seq libraries were prepared according to manufacturer’s instructions using the Illumina TruSeq Stranded mRNA Kit and 50 bp single reads sequencing was performed with the Illumina HiSeq 4000 Sequencer. For the smaller amounts of RNA obtained from cardiomyocyte nuclei (n = 3 for each mouse strain), RNA-seq library preparation was performed using the SMARTer Stranded Total RNA-Seq Kit v2 - Pico (Takara Bio, Mountain View, CA) prior to 100 bp paired-end sequencing. For the last two sequencing analyses, after assessing the read quality using FastQC 0.11.8, alignment on the mouse GRCm38/mm10 genome was performed using STAR 2.5.1b. For all RNA-seq data, feature quantification was extracted with featureCounts v1.6.0 on the GRCm38 v98 Ensembl annotation. We also quantified additional transcript in the *Uty* region (AK142197, AK153586, Gm39552 and C030026M15) as there were not included in the Ensembl reference annotation. Differential expression analysis was performed with DESeq2 1.22.2 R package. For the nucRNA-Seq analysis, additional steps were taken to exclude artifactual contamination with genomic DNA. In particular, we identified a total of 2 megabases of regions where the absence of transcription was assessed on the basis of the following two overlapping criteria: *i*) lack of data showing expression of corresponding transcripts in hearts (on the basis of mouse ENCODE transcriptome data), in contrast to other non-cardiac tissues where the pattern of tissue expression was restricted to just a few tissue types; and *ii*) lack of data for chromatin accessibility in the same regions, on the basis of ENCODE Dnase hypersensitivity tracks obtained in mouse adult hearts. UCSC exons and introns of these regions were extracted with GenomicFeatures R package and quantified as FPKM values with featureCounts. Absence of genomic contamination was confirmed by verifying that FPKM values were either zero or very low (Supplemental Table S6).

ChIP-Seq was performed on two replicate samples from each mouse strain as we described previously ^54^, using the ChIP grade ab4729 anti-H3K27 antibody from Abcam (Cambridge, MA). For the analysis, read quality was confirmed using FastQC 0.11.8 before alignment using Bowtie2 2.2.6 on the mouse GRCm37/mm9 genome (that latter build was chosen in order to allow for comparisons with other ChIP tracks published by ENCODE). Peak calling and differential enrichment were performed using the MACS2 2.0.10 “callpeaks” and “bdgdiff” functions, respectively. For ATAC-Seq, preparations containing 50,000 nuclei from either WT or *Uty^GT^* hearts were performed on three different days, and processed as described previously^55^. For the analysis, sequences from the adapters (NexteraPE-PE) were removed with Trimmomatic 0.36 after read quality control. Alignment was performed as for ChIP-Seq post-processing to remove PCR duplicates (Picard tool 2.4.1) and reads mapping to mitochondrial DNA (samtools 1.8). In order to represent the real Tn5 transposase binding sites of 9bp, the coordinates of the reads were shifted by 4bp for the plus strand and by 5bp for the minus strand, using Deeptools 3.0.1. The former was also used to remove ENCODE’s blacklisted regions (signal artefact regions) and convert Bam files to BEDPE format. MACS2 was used to identify significant peaks using q-value < 0.01. Diffbind 2.10.0 R package was used to generate the count matrix of Tn5 insertion site numbers for each consensus peak (peaks that were present in at least 2 samples). Differentially accessibility regions (DARs) were identified between conditions using edgeR R package 3.24.3, including its generalized linear model functionalities to perform a multi-factor analysis to control for batch effects.

### Data deposition

Results from all the sequencing-based experiments were deposited in the Gene Expression Omnibus database as SuperSeries GSE147652.

### Molecular biology

Full-length mUty cDNA was obtained by PCR amplification of reverse-transcribed mouse heart RNA, then subcloned into the pIRES-V5 vector (gift of N. Seidah). After full sequence verification, the whole cDNA was subcloned into the lentiviral CMV GFP destination vector 736-1 (Addgene #19732), and further used along with packaging vectors to generate lentiviral particles (control lentiviral particles were generated in the same manner using empty lentiviral vectors). For infection, confluent P1 mouse fibroblasts were passaged at a concentration of 50,000 cells / 2 cm^2^ wells. Twenty four hours after replating, the cells were exposed for 6 hours to lentiviral particles (M.O.I. = 3) in growth medium supplemented with 6 μg/ml dextran sulfate, then washed and grown further overnight. Cell were then trypsinized for cytofluorometric separation of GFP-positive (i.e. cells with lentiviral integration) and GFP-negative cells. A total of 40-60,000 cells were then used for each sample to extract total RNA using the RNeasy micro RNA kit.

RT-qPCR was performed on cDNA obtained from total RNA using the High-Capacity cDNA Reverse Transcription Kit (ThermoFisher), using the primers detailed in Supplementary Table S3. Results were expressed as relative expression ratios of the 2(-ΔΔCt) values of the gene of interest vs. of that of the *Rps16* normalizing gene.

### Western blotting

After separation of gels and transfer to PVDF membranes, the latter were incubated with the following primary antibodies: monoclonal anti DDX3 C4, raised against the 114 N-terminal portion of human DDX3x (Santa Cruz Biotehcnology); polyclonal anti DDX3, raised against recombinant human DDX3x (Genetex, Irvine, CA); polyclonal anti GAPDH (Proteintech, Rosemont, IL). After treatment with horse radish peroxidase coupled second antibodies, signal was revealed by chemiluminescence, then visualized and analyzed using the Image lab software (Bio-Rad Laboratories, Saint-Laurent, QC, Canada).

## SUPPLEMENTARY FILES

### Supplementary tables

**Supplementary Table S1:** List of mouse genes with highest levels of co-expression with Uty, ranked on the basis of mutual ranking (MR) scores

| Rank | Gene | Chr | Gene type | Entrez Gene ID | MR |
| --- | --- | --- | --- | --- | --- |
| 0 | Uty | chrY:1,097,144-1,245,718 | protein coding | 22290 | 0 |
| 1 | Ddx3y | chrY:1,260,771-1,286,629 | protein coding | 26900 | 2.13 |
| 2 | Eif2s3y | chrY:1,010,543-1,028,594 | protein coding | 26908 | 2.26 |
| 3 | Kdm5d | chrY:897,790-946,316 | protein coding | 20592 | 2.28 |
| 4 | Gm39552 | chrY:1,114,067-1,120,066 | lncRNA | 105243746 | 25.03 |
| 5 | C030026M15Rik | chrY:1,161,073-1,162,805 | lncRNA | 77378 | 116.33 |
| 6 | Uba1y | chrY:2,264,257-2,278,687 | protein coding | 22202 | 218.38 |
| 7 | Gm35612 | chrX:50,537,389-50,562,598 | lncRNA | 102639259 | 750.5 |
| 8 | Usp9y | chrY:1,298,961-1,459,782 | protein coding | 107868 | 1590.67 |
| 9 | 4932431L22Rik | chrY:70,367,444-70,370,583 | lncRNA | 619557 | 2289.96 |

Co-expression values are expressed as “mutual ranking scores”, which corresponded to the mean geometric mean of the correlation rank of gene A to gene B and of gene B to gene A. Calculations were performed with tools provided by COEXPRESdb, using data from 2236 Affymetrix arrays (microarray platform GPL1261) performed on multiple tissues across 154 experiments. A detailed list of all tissues used is available at the following link: (https://coexpresdb.jp/static/help/GPL1261v2_samples.shtml). The MR scores are inversely proportional to the strength of co-expression levels (the lower the score, the stronger the interaction). MR scores higher than 200 are considered as weak.

**Supplementary Table S2:** Top most differentially accessible regions in chromatin from cardiomyocyte nuclei from either WT of *Uty^GT^* mice, as determined by ATAC-Seq assays. Regions formatted in bold correspond to those identified in the context of other experiments. Regions with FDR significance < E-09 are shown only if they are flanked by other regions with some levels of significance.

| Gene Name | Chr | Start | End |  | FDR |
| --- | --- | --- | --- | --- | --- |
| **Uty** | **chrY** | **581836** | **582382** | **Promoter-TSS** | **3.50E-34** |
| En2 | chr5 | 28495275 | 28496134 | First intron | 1.02E-31 |
| **Ddx3y** | **chrY** | **622332** | **623154** | **First intron** | **2.86E-18** |
| **Eif2s3y** | **chrY** | **346529** | **347077** | **Promoter-TSS** | **5.50E-14** |
| **Malat1** | **chr19** | **5821944** | **5823927** | **Intergenic** | **9.64E-19** |
| Neat1 | chr19 | 5824319 | 5825012 | TTS | 8.61E-13 |
| Neat1 | chr19 | 5831054 | 5832841 | Non-coding | 2.23E-09 |
| Neat1 | chr19 | 5838441 | 5842512 | Non-coding | 8.52E-09 |
| Neat1 | chr19 | 5828977 | 5829566 | Non-coding | 1.12E-08 |
| Neat1 | chr19 | 5829981 | 5830325 | Non-coding | 3.04E-07 |
| Neat1 | chr19 | 5837107 | 5838143 | Non-coding | 1.43E-06 |
| Neat1 | chr19 | 5835340 | 5836037 | Non-coding | 2.01E-06 |
| Neat1 | chr19 | 5833417 | 5833811 | Non-coding | 2.73E-04 |
| Myh7 | chr14 | 55619384 | 55620431 | Promoter-TSS | 1.60E-09 |
| Myh7 | chr14 | 55620990 | 55624433 | Intergenic | 1.16E-06 |
| Myh7 | chr14 | 55612681 | 55615300 | Intergenic | 2.74E-06 |
| Myh7 | chr14 | 55608473 | 55609053 | Intron 12 of 40 | 4.91E-05 |
| Myh7 | chr14 | 55615784 | 55616173 | Intergenic | 1.78E-03 |
| Myh7 | chr14 | 55606934 | 55608000 | Intron 15 of 40 | 1.86E-02 |
| Zbtb16 | chr9 | 48632529 | 48633073 |  | 5.69E-09 |
| Zbtb16 | chr9 | 48574572 | 48575021 |  | 1.91E-05 |
| Zbtb16 | chr9 | 48564539 | 48564746 |  | 2.63E-04 |
| Zbtb16 | chr9 | 48590823 | 48591848 |  | 4.93E-04 |
| Zbtb16 | chr9 | 48605695 | 48605938 |  | 7.61E-04 |
| Zbtb16 | chr9 | 48568245 | 48568945 |  | 2.21E-02 |
| Zbtb16 | chr9 | 48636958 | 48637403 |  | 2.73E-02 |
| Zbtb16 | chr9 | 48622674 | 48623269 |  | 4.43E-02 |
| Plin4 | chr17 | 56242595 | 56246147 |  | 2.13E-08 |
| Plin4 | chr17 | 56239471 | 56241031 |  | 1.88E-06 |

**Supplementary Table S3:**
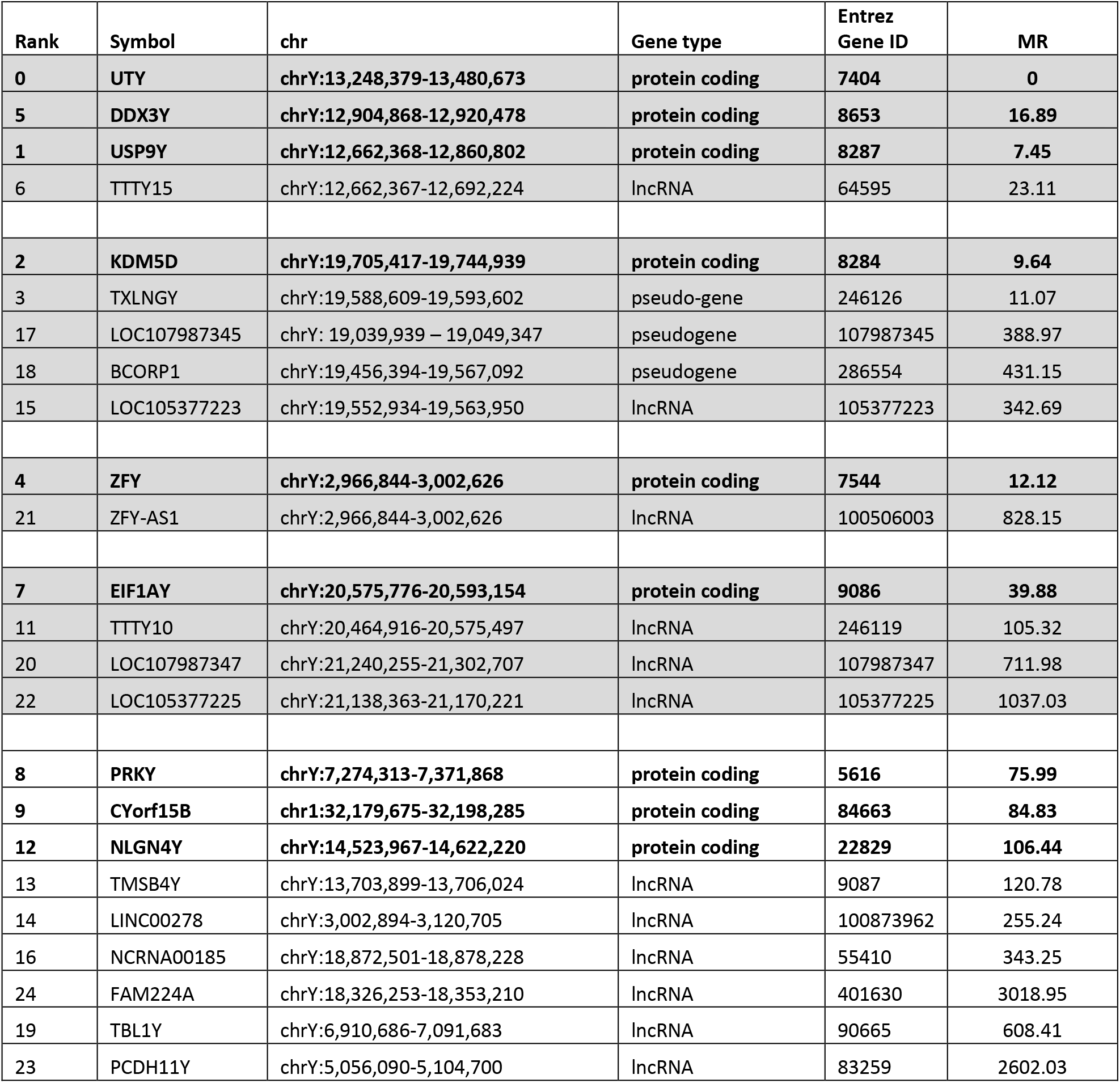
List of human genes with highest levels of co-expression with Uty, as determined on the basis of mutual ranking (MR) scores. Genes are shown in shaded cells when other neighboring genes also show evidence of interaction with UTY.

**Supplementary Table S4:** Coordinates of genomic regions predicted to be transcriptionally silent in cardiomyocytes. The negligeable FPKM values obtained for these regions in samples from either *Y^UtyGT^* mice or WT mice (identified as either Uty* or b6*) were interpreted as evidence for the absence of DNA contaminationin the nuclear RNA samples.

| ensembl_gene_id | Coordinates mm10 | length (bp) | b6_11 | b6_15 | b6_31 | uty_11 | uty_15 | uty_31 |
| --- | --- | --- | --- | --- | --- | --- | --- | --- |
| ENSMUSG00000008789 | chr7:17,713,252-17,761,121 | 47,870 bp | 0.00 | 0.00 | 0.00 | 0.00 | 0.00 | 0.00 |
| ENSMUSG00000019785 | chr10:33,512,334-33,624,600 | 112,267 bp | 0.00 | 0.00 | 0.00 | 0.00 | 0.00 | 0.00 |
| ENSMUSG00000020633 | chr12:28,437,795-28,548,337 | 110,543 bp | 0.00 | 0.00 | 0.00 | 0.00 | 0.00 | 0.00 |
| ENSMUSG00000000560 | chr5:70,961,057-71,095,849 | 134,793 bp | 0.04 | 0.07 | 0.13 | 0.10 | 0.11 | 0.06 |
| ENSMUSG00000001260 | chr5:70,751,047-70,842,617 | 91,571 bp | 0.00 | 0.00 | 0.00 | 0.00 | 0.00 | 0.00 |
| ENSMUSG00000018589 | chrX:165,129,017-165,326,981 | 197,965 bp | 0.00 | 0.00 | 0.00 | 0.00 | 0.00 | 0.00 |
| ENSMUSG00000022112 | chr14:115,092,215-116,525,192 | 1,432,978 bp | 0.00 | 0.00 | 0.00 | 0.00 | 0.00 | 0.00 |
| ENSMUSG00000017688 | chr3:3,508,030-3,659,800 | 151,771 bp | 0.00 | 0.00 | 0.00 | 0.00 | 0.00 | 0.00 |
| ENSMUSG00000022076 | chr14:96,105,265-96,519,034 | 413,770 bp | 0.00 | 0.00 | 0.00 | 0.00 | 0.00 | 0.00 |
| ENSMUSG00000009356 | chr11:87,806,428-87,826,114 | 19,687 bp. | 0.00 | 0.00 | 0.00 | 0.00 | 0.00 | 0.00 |
| ENSMUSG00000021363 | chr13:41,025,120-41,079,706 | 54,587 bp | 0.00 | 0.00 | 0.00 | 0.00 | 0.00 | 0.01 |
| ENSMUSG00000005493 | chr3:153,857,141-153,906,138 | 48,998 bp | 0.18 | 0.10 | 0.15 | 0.22 | 0.10 | 0.22 |
| ENSMUSG00000010505 | chr2:181,763,332-181,827,775 | 64,444 bp | 0.00 | 0.00 | 0.00 | 0.00 | 0.00 | 0.00 |
| ENSMUSG00000004231 | chr19:44,757,394-44,837,871 | 80,478 bp | 0.00 | 0.00 | 0.00 | 0.00 | 0.01 | 0.00 |
| ENSMUSG00000021852 | chr14:49,298,520-49,525,837 | 227,318 bp | 0.00 | 0.00 | 0.00 | 0.01 | 0.01 | 0.00 |
| ENSMUSG00000010435 | chr8:12,573,049-12,600,738 | 27,690 bp | 0.00 | 0.00 | 0.00 | 0.00 | 0.00 | 0.00 |
| ENSMUSG00000003271 | chr7:45,729,983-45,759,555 | 29,573 bp | 0.00 | 0.01 | 0.00 | 0.00 | 0.00 | 0.00 |
| ENSMUSG00000001670 | chr8:109,990,436-109,999,804 | 9,369 bp | 0.00 | 0.00 | 0.00 | 0.00 | 0.00 | 0.00 |
| ENSMUSG00000021541 | chr13:56,773,098-56,895,789 | 122,692 bp | 0.00 | 0.00 | 0.00 | 0.00 | 0.00 | 0.03 |

**Supplementary Table S5:**
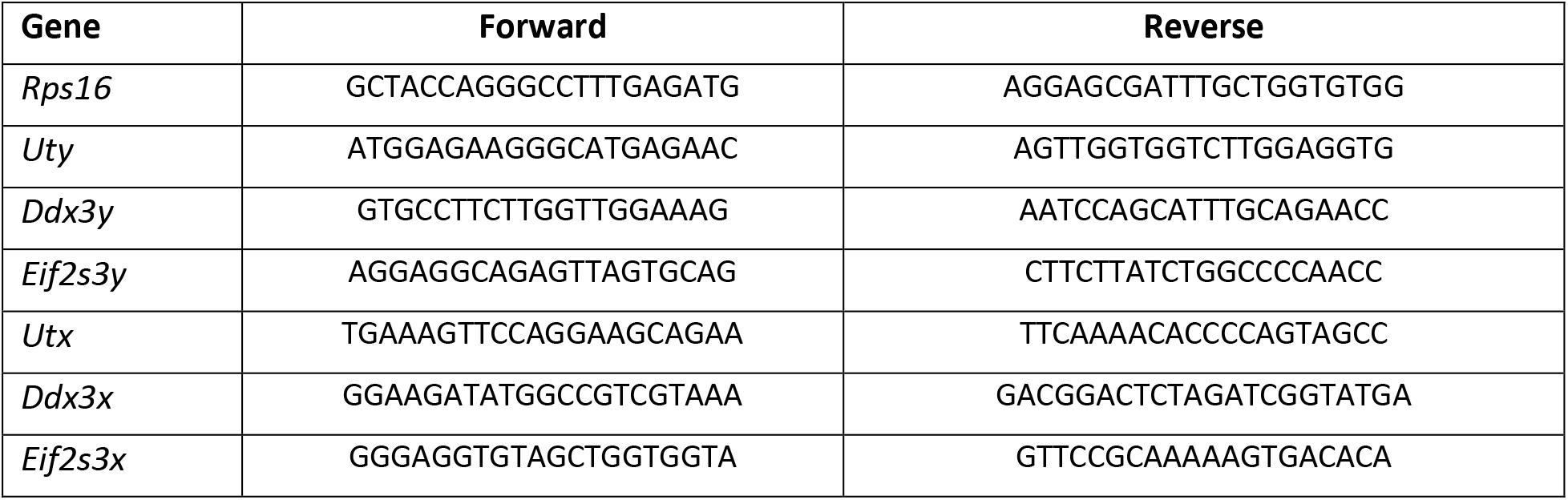
List of oligomer primers used for RT-qPCR amplification

**<u>Supplementary Table S6</u>** is provided in an additional file as a four tab-containing Excel spreadsheet. Each sheet lists all GO terms showing both high (> 2 fold) and significant (FDR < 0.01) enrichment among genes affected by *Uty^GT^* (either down-or upregulated) in either left ventricles (LV) or cardiomyocyocytes (CM).

**<u>Supplementary Table S7</u>** is provided in an additional file as a four tab-containing Excel spreadsheet. It contains information about genes showing either up- or down-regulation in left ventricles from *Uty^GT^* mice, as well as a more comprehensive list of GO categories showing significant enrichment for these genes.

**<u>Supplementary Table S8</u>** is provided in an additional file as a four tab-containing Excel spreadsheet. It contains information about genes showing either up- or down-regulation in cardiomyocytes from *Uty^GT^* mice, as well as a more comprehensive list of GO categories showing significant enrichment for these genes.

### Supplementary figures

**Supplementary Fig. S1:**
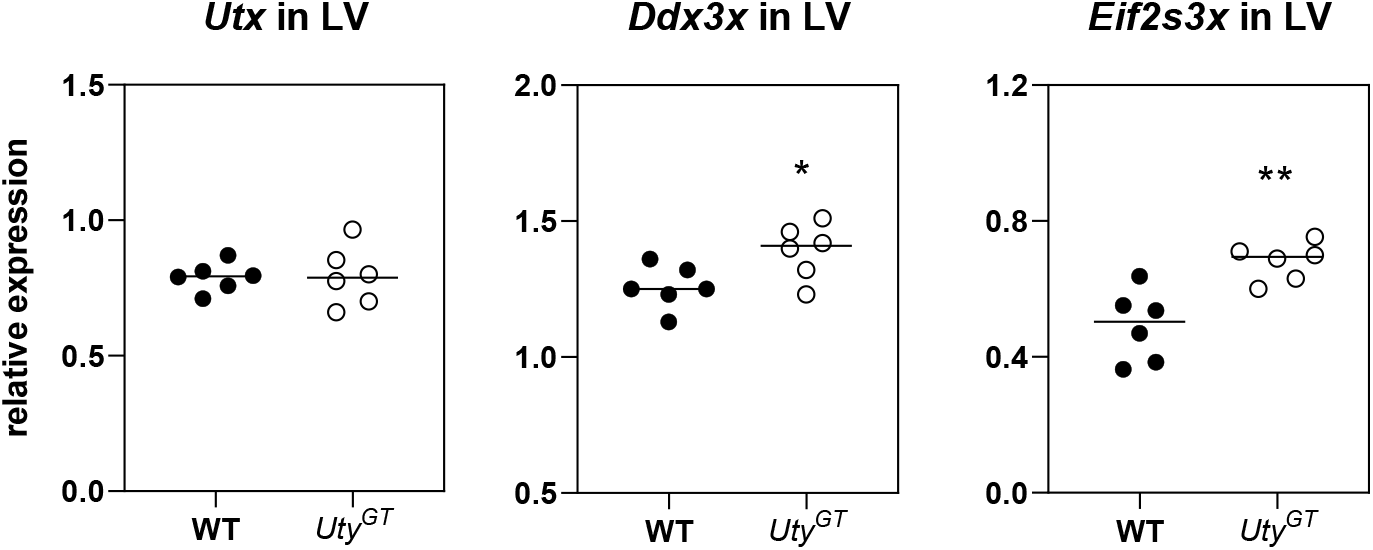
RT-qPCR quantification of abundance of mRNA transcripts of *Utx, Ddx3x and Eif2s3x in left ventricular extracts from Uty^GT^ and WT mice*. Expression of *Utx* was unaffected, *Ddx3x* showed a slight increase of about 10 % (\**P* < 0.05), expression of *Eif2s3x* was increased by about 40% (*P* < 0.01).

**Supplementary Fig. S2:**
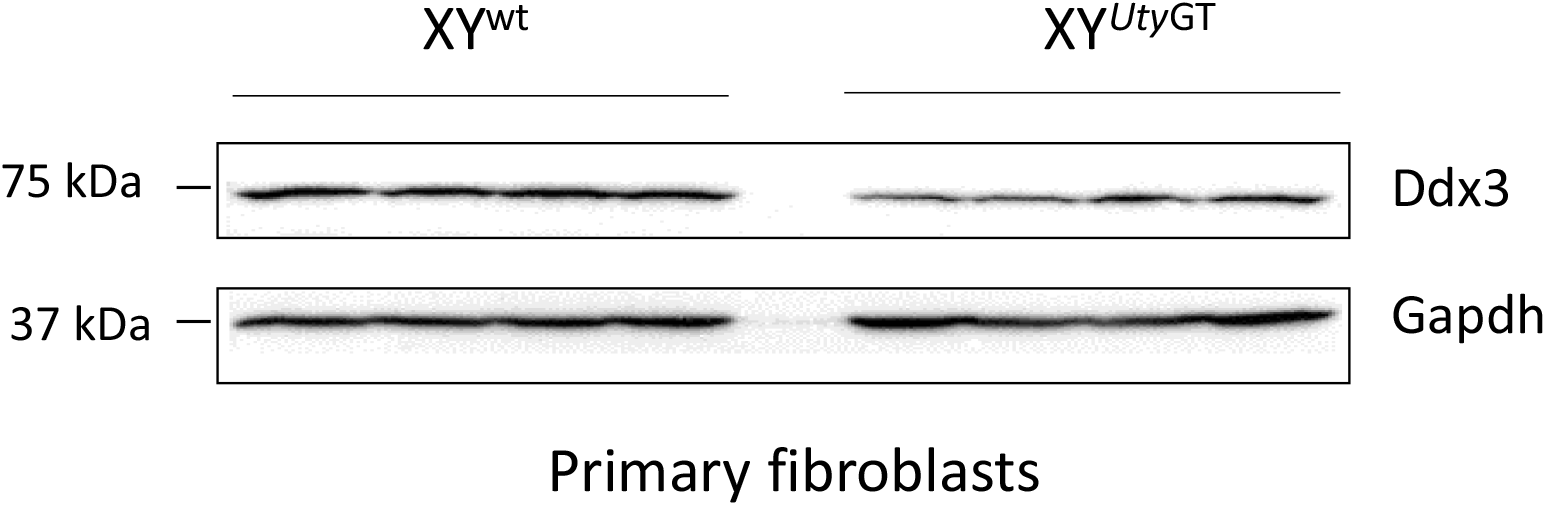
Western blot analysis of primary fibroblasts from either *Uty^GT^ and WT mice*. Upper part: signal obtained with anti-DDX3 antibodies from Genetex; bottom part: signal obtained with anti-GAPDH antibodies.

**Supplementary Fig. S3:**
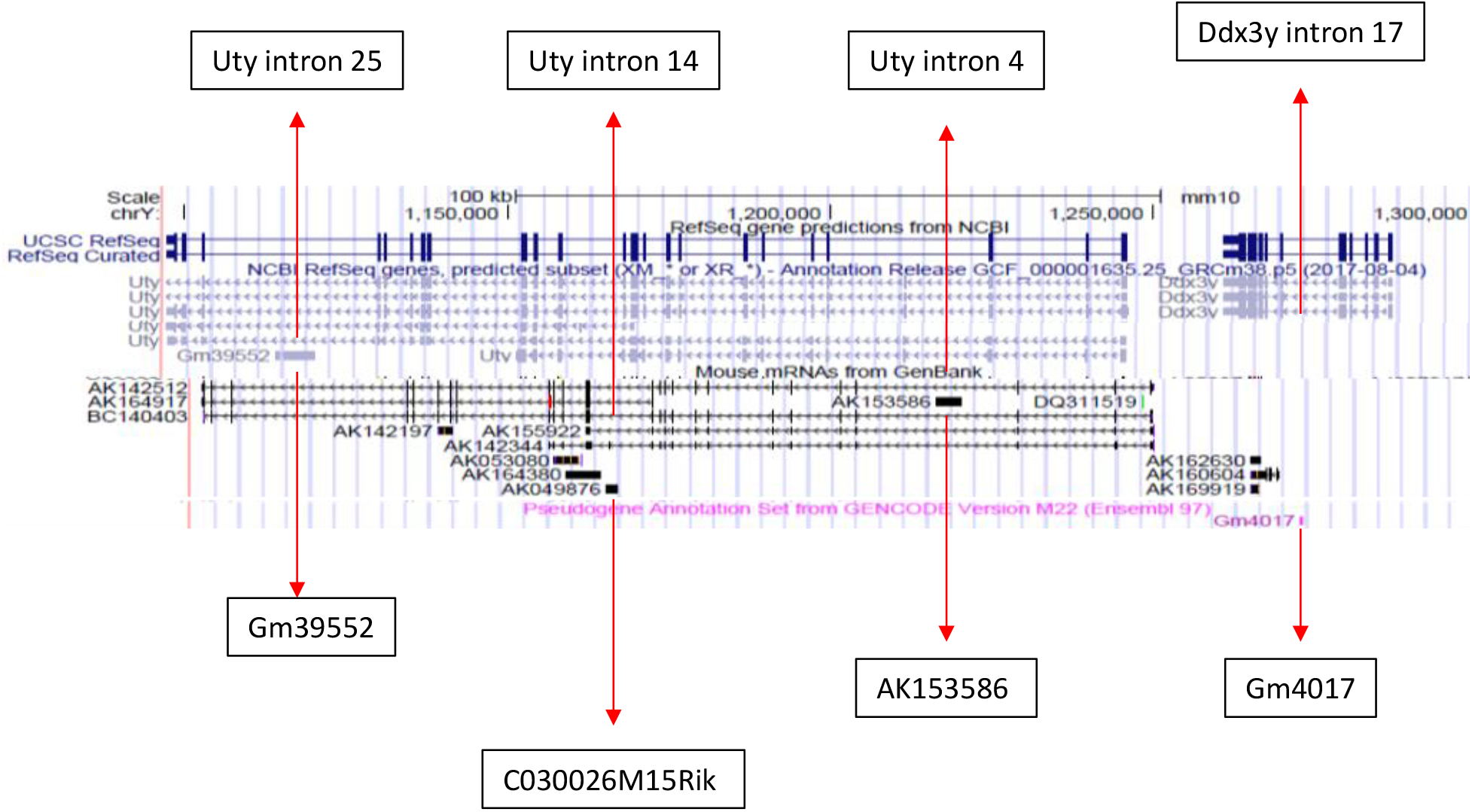
Localization of loci embedded within introns of either Uty or Ddx3y and containing sequences corresponding to putatively expressed transcripts.

**Supplementary Fig. S4:**
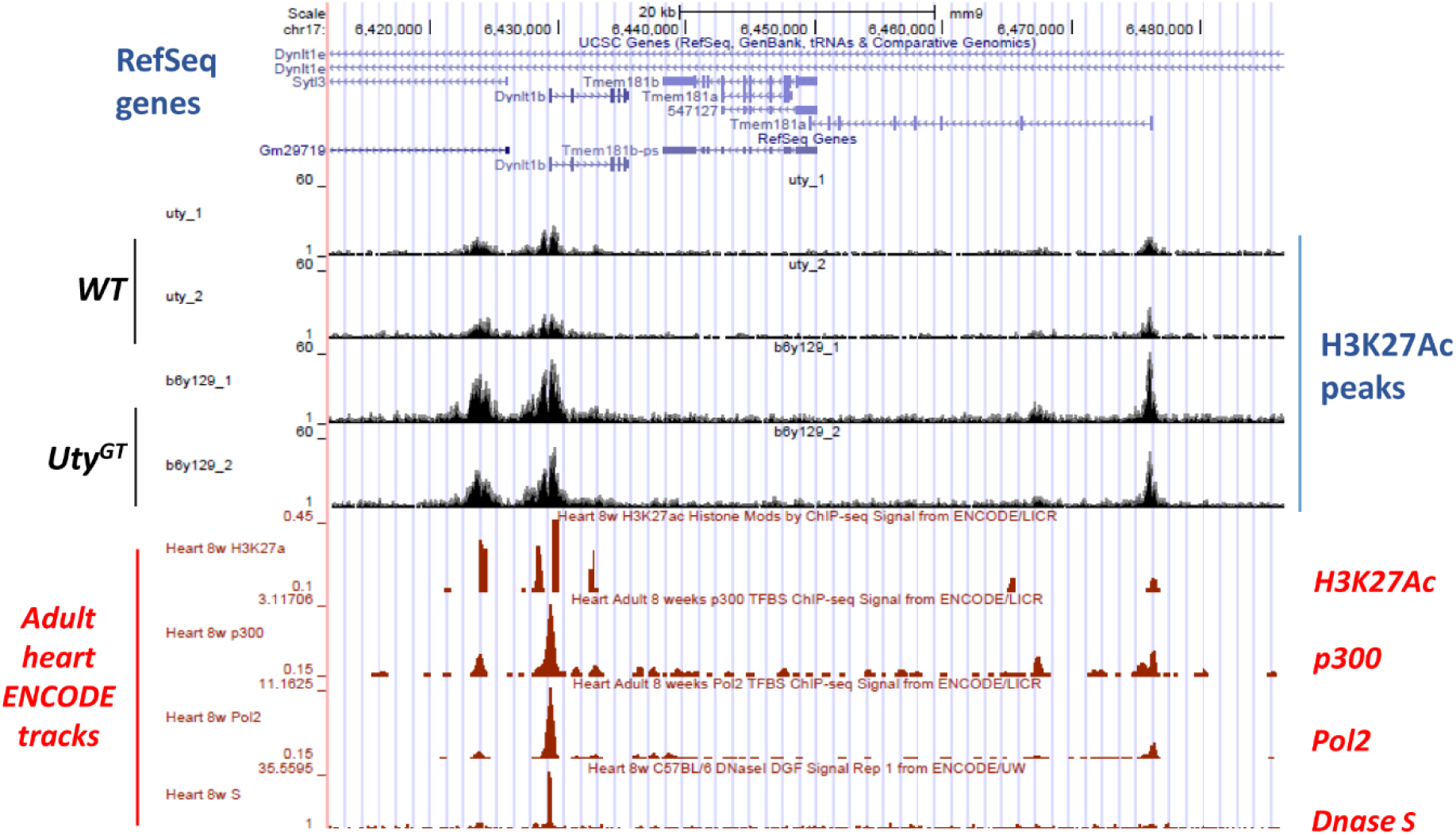
Graphic representation of ATAC-Seq results for chr17. Regions shown contain peaks showing significant differences in intensity between *Uty^GT^* and WT mice. For comparison, ENCODE tracks obtained using extracts from adult mouse hearts for either H3K27ac, p300 or Pol2 ChIP-Seq or Dnase hypersensitivity assays are also shown (in red). Loci of known genes are shown in the top part of the figure (in blue).

**Supplementary Fig. S5:**
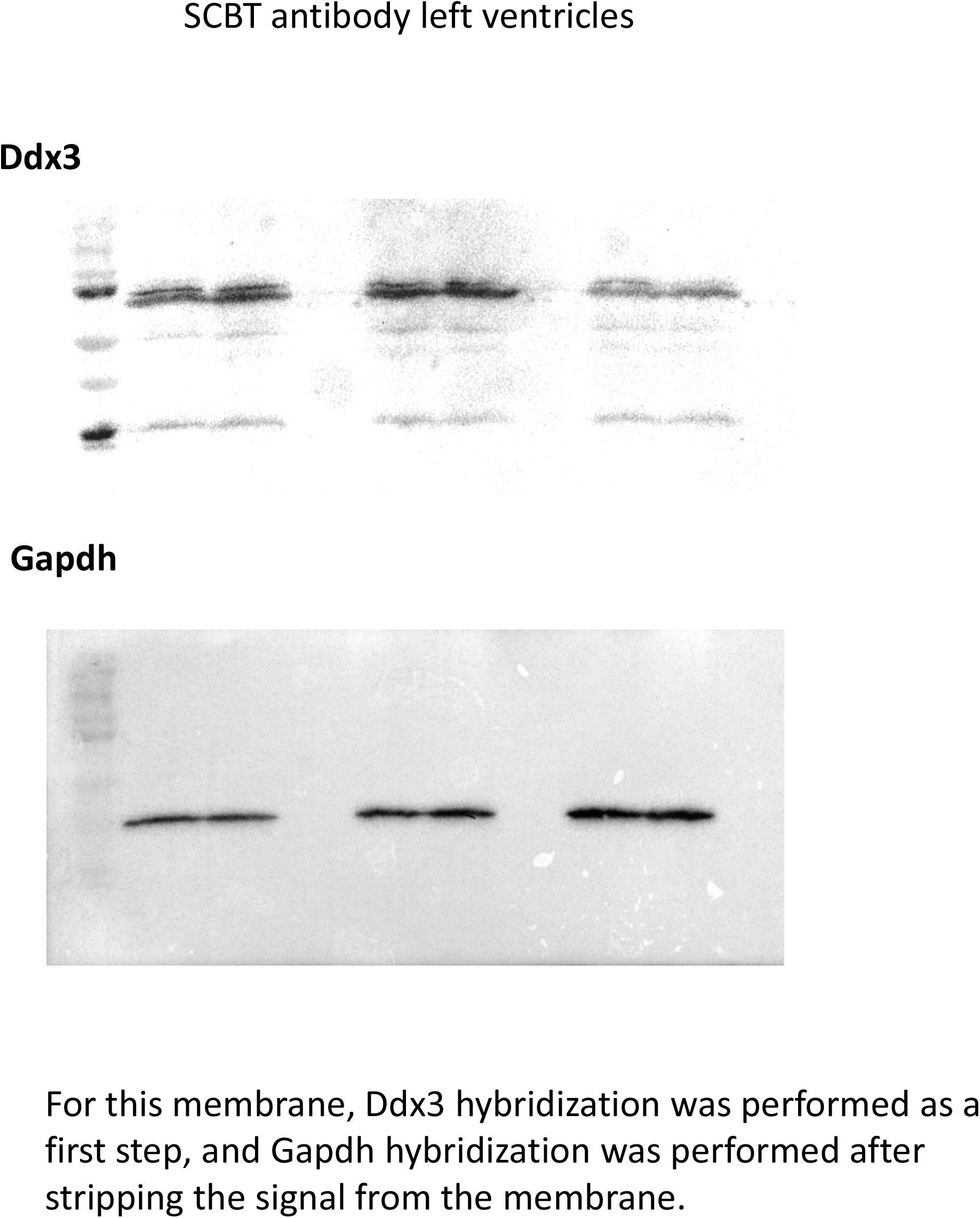
Full images of the hybridized membranes that have been used for the top part of Fig. 2 in the manuscript.

**Supplementary Fig. S6:**
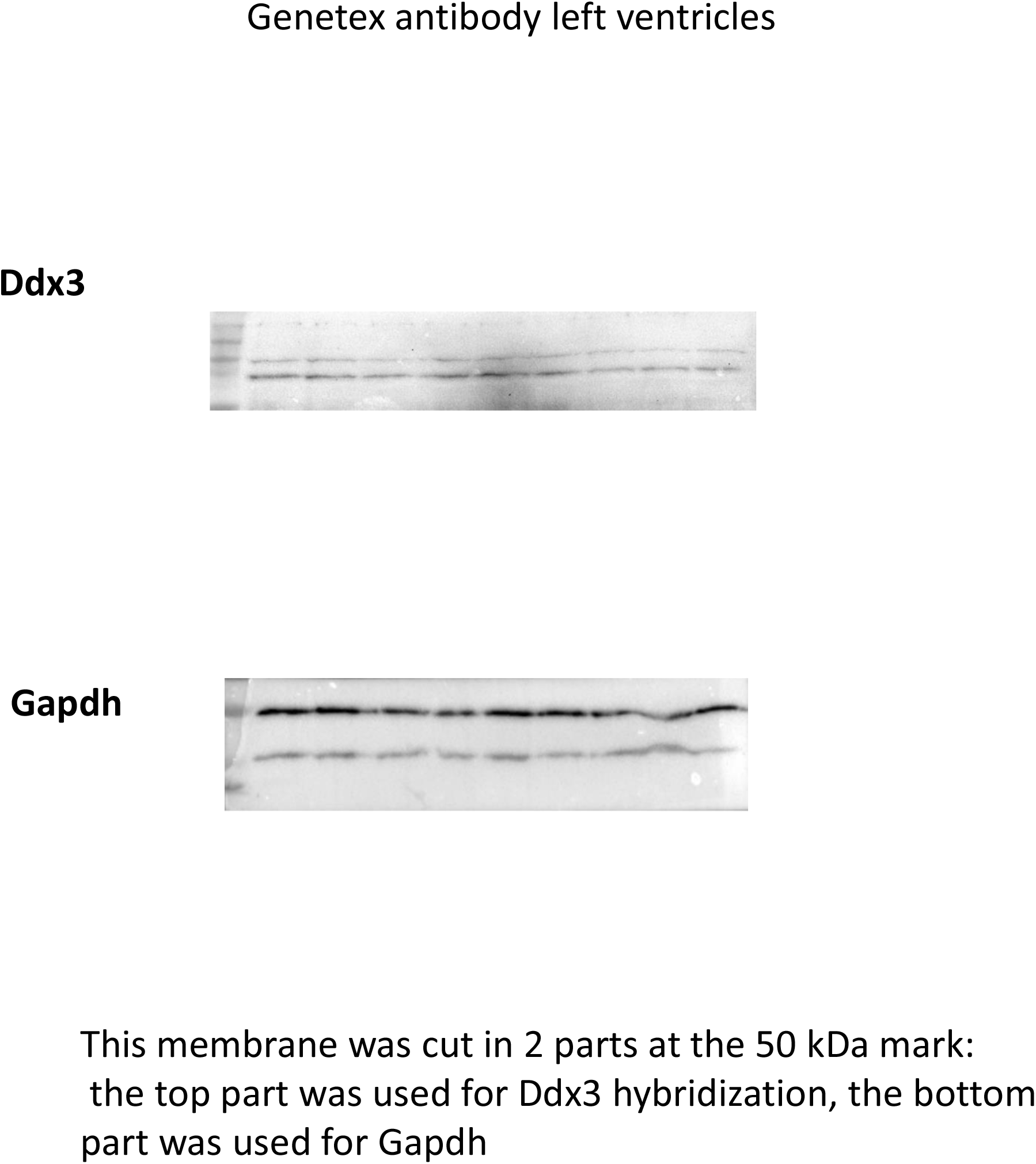
Full images of the hybridized membranes that have been used for the bottom part of Fig. 2 in the manuscript.

## REFERENCES

1. Prokop, J. W. & Deschepper, C. F. Chromosome Y genetic variants: impact in animal models and on human disease. Physiological Genomics 47, 525–537 (2015).

2. Case, L. K. & Teuscher, C. Y genetic variation and phenotypic diversity in health and disease. Biol Sex Differ 6, (2015).

3. Krementsov, D. N. et al. Genetic variation in chromosome Y regulates susceptibility to influenza A virus infection. PNAS 114, 3491–3496 (2017).

4. Prokop, J. W. et al. The phenotypic impact of the male-specific region of chromosome-Y in inbred mating: the role of genetic variants and gene duplications in multiple inbred rat strains. Biology of Sex Differences 7, 10 (2016).

5. AlSiraj, Y. et al. XX sex chromosome complement promotes atherosclerosis in mice. Nat Commun 10, 1–13 (2019).

6. Eales James M. et al. Human Y Chromosome Exerts Pleiotropic Effects on Susceptibility to Atherosclerosis. Arteriosclerosis, Thrombosis, and Vascular Biology 0, ATVBAHA.119.312405.

7. Forsberg, L. A. et al. Mosaic loss of chromosome Y in peripheral blood is associated with shorter survival and higher risk of cancer. Nat Genet 46, 624–628 (2014).

8. Dumanski, J. P. et al. Mosaic Loss of Chromosome Y in Blood Is Associated with Alzheimer Disease. The American Journal of Human Genetics 98, 1208–1219 (2016).

9. Grassmann, F. et al. Y chromosome mosaicism is associated with age-related macular degeneration. Eur J Hum Genet 27, 36–41 (2019).

10. Haitjema, S. et al. Loss of Y Chromosome in Blood Is Associated With Major Cardiovascular Events During Follow-Up in Men After Carotid Endarterectomy. Circulation: Cardiovascular Genetics (2017).

11. Skaletsky, H. et al. The male-specific region of the human Y chromosome is a mosaic of discrete sequence classes. Nature 423, 825–837 (2003).

12. Bellott, D. W. et al. Mammalian Y chromosomes retain widely expressed dosage-sensitive regulators. Nature 508, 494–499 (2014).

13. Sekiguchi, T., Iida, H., Fukumura, J. & Nishimoto, T. Human DDX3Y, the Y-encoded isoform of RNA helicase DDX3, rescues a hamster temperature-sensitive ET24 mutant cell line with a DDX3X mutation. Experimental Cell Research 300, 213–222 (2004).

14. Wang, T. et al. Identification and characterization of essential genes in the human genome. Science 350, 1096–1101 (2015).

15. Shpargel, K. B., Sengoku, T., Yokoyama, S. & Magnuson, T. UTX and UTY Demonstrate Histone Demethylase-Independent Function in Mouse Embryonic Development. PLoS Genet 8, e1002964 (2012).

16. Gozdecka, M. et al. UTX-mediated enhancer and chromatin remodeling suppresses myeloid leukemogenesis through noncatalytic inverse regulation of ETS and GATA programs. Nature Genetics 50, 883–894 (2018).

17. Gažová, I., Lengeling, A. & Summers, K. M. Lysine demethylases KDM6A and UTY: The X and Y of histone demethylation. Molecular Genetics and Metabolism (2019) doi:10.1016/j.ymgme.2019.04.012.

18. Maan, A. A. et al. The Y chromosome: a blueprint for men’s health? Eur J Hum Genet 25, 1181–1188 (2017).

19. Walport, L. J. et al. Human UTY(KDM6C) Is a Male-specific N∈-Methyl Lysyl Demethylase. J. Biol. Chem. 289, 18302–18313 (2014).

20. Van der Meulen, J., Speleman, F. & Vlierberghe, P. V. The H3K27me3 demethylase UTX in normal development and disease. Epigenetics 9, 658–668 (2014).

21. Wang, S.-P. et al. A UTX–MLL4–p300 Transcriptional Regulatory Network Coordinately Shapes Active Enhancer Landscapes for Eliciting Transcription. Mol Cell 67, 308–321.e6 (2017).

22. Ford, D. J. & Dingwall, A. K. The cancer COMPASS: navigating the functions of MLL complexes in cancer. Cancer Genetics 208, 178–191 (2015).

23. Obayashi, T. & Kinoshita, K. COXPRESdb: a database to compare gene coexpression in seven model animals. Nucleic Acids Res. 39, D1016–1022 (2011).

24. Mitchell, J. A. et al. Nuclear RNA Sequencing of the Mouse Erythroid Cell Transcriptome. PLOS ONE 7, e49274 (2012).

25. Preissl, S. et al. Deciphering the Epigenetic Code of Cardiac Myocyte TranscriptionNovelty and Significance. Circulation Research 117, 413–423 (2015).

26. Li, G. et al. Comparative analysis of mammalian Y chromosomes illuminates ancestral structure and lineage-specific evolution. Genome Res. 23, 1486–1495 (2013).

27. Irimia, M. et al. Extensive conservation of ancient microsynteny across metazoans due to cis-regulatory constraints. Genome Res. 22, 2356–2367 (2012).

28. Scott-Boyer, M.-P. & Deschepper, C. F. Genome-Wide Detection of Gene Coexpression Domains Showing Linkage to Regions Enriched with Polymorphic Retrotransposons in Recombinant Inbred Mouse Strains. G3 3, 597–605 (2013).

29. Bellott, D. W. & Page, D. C. Reconstructing the Evolution of Vertebrate Sex Chromosomes. Cold Spring Harb Symp Quant Biol 74, 345–353 (2009).

30. Sun, Q., Hao, Q. & Prasanth, K. V. Nuclear long noncoding RNAs: key regulators of gene expression. Trends Genet 34, 142–157 (2018).

31. Ditton, H. J., Zimmer, J., Kamp, C., Meyts, E. R.-D. & Vogt, P. H. The AZFa gene DBY (DDX3Y) is widely transcribed but the protein is limited to the male germ cells by translation control. Hum. Mol. Genet. 13, 2333–2341 (2004).

32. Jaroszynski, L. et al. Translational control of the AZFa gene DDX3Y by 5’UTR exon-T extension. International Journal of Andrology 34, 313–326 (2011).

33. Rosinski, K. V. et al. DDX3Y encodes a class I MHC–restricted H-Y antigen that is expressed in leukemic stem cells. Blood 111, 4817–4826 (2008).

34. Zorn, E. et al. Minor Histocompatibility Antigen DBY Elicits a Coordinated B and T Cell Response after Allogeneic Stem Cell Transplantation. Journal of Experimental Medicine 199, 1133–1142 (2004).

35. Vakilian, H. et al. DDX3Y, a Male-Specific Region of Y Chromosome Gene, May Modulate Neuronal Differentiation. J. Proteome Res. 14, 3474–3483 (2015).

36. Szappanos, D. et al. The RNA helicase DDX3X is an essential mediator of innate antimicrobial immunity. PLOS Pathogens 14, e1007397 (2018).

37. Khramtsova, E. A., Davis, L. K. & Stranger, B. E. The role of sex in the genomics of human complex traits. Nature Reviews Genetics 20, 173 (2019).

38. Mayne, B. T. et al. Large Scale Gene Expression Meta-Analysis Reveals Tissue-Specific, Sex-Biased Gene Expression in Humans. Front. Genet. 7, (2016).

39. Ariumi, Y. Multiple functions of DDX3 RNA helicase in gene regulation, tumorigenesis, and viral infection. Front. Genet. 5, 423 (2014).

40. Sharma, D. & Jankowsky, E. The Ded1/DDX3 subfamily of DEAD-box RNA helicases. Critical Reviews in Biochemistry and Molecular Biology 49, 343–360 (2014).

41. Lee, C.-S. et al. Human DDX3 functions in translation and interacts with the translation initiation factor eIF3. Nucleic Acids Res 36, 4708–4718 (2008).

42. Soto-Rifo, R. et al. DEAD-box protein DDX3 associates with eIF4F to promote translation of selected mRNAs. The EMBO Journal 31, 3745–3756 (2012).

43. Geissler, R., Golbik, R. P. & Behrens, S.-E. The DEAD-box helicase DDX3 supports the assembly of functional 80S ribosomes. Nucleic Acids Res 40, 4998–5011 (2012).

44. Fazi, F. & Fatica, A. Interplay Between N6-Methyladenosine (m6A) and Non-coding RNAs in Cell Development and Cancer. Front. Cell Dev. Biol. 7, (2019).

45. Leppek, K., Das, R. & Barna, M. Functional 5’ UTR mRNA structures in eukaryotic translation regulation and how to find them. Nat Rev Mol Cell Biol 19, 158–174 (2018).

46. Lei, L. et al. Functions and regulatory mechanisms of metastasis-associated lung adenocarcinoma transcript 1. Journal of Cellular Physiology 234, 134–151 (2019).

47. Shah, A. et al. The DEAD-Box RNA Helicase DDX3 Interacts with m6A RNA Demethylase ALKBH5. Stem Cells International https://www.hindawi.com/journals/sci/2017/8596135/ (2017) doi:https://doi.org/10.1155/2017/8596135.

48. Arnold, A. P. & Lusis, A. J. Understanding the Sexome: Measuring and Reporting Sex Differences in Gene Systems. Endocrinology 153, 2551–2555 (2012).

49. Seluanov, A., Vaidya, A. & Gorbunova, V. Establishing Primary Adult Fibroblast Cultures From Rodents. J Vis Exp (2010) doi:10.3791/2033.

50. O’Connell, T. D., Rodrigo, M. C. & Simpson, P. C. Isolation and culture of adult mouse cardiac myocytes. Methods Mol. Biol. 357, 271–296 (2007).

51. Bergmann, O. & Jovinge, S. Isolation of Cardiomyocyte Nuclei from Post-mortem Tissue. e4205 (2012) doi:10.3791/4205.

52. Stone, C. M. et al. Inhibition of homologous phosphorolytic ribonucleases by citrate may represent an evolutionarily conserved communicative link between RNA degradation and central metabolism. Nucleic Acids Res 45, 4655–4666 (2017).

53. Hancock, R. & Hadj-Sahraoui, Y. Isolation of Cell Nuclei Using Inert Macromolecules to Mimic the Crowded Cytoplasm. PLOS ONE 4, e7560 (2009).

54. Praktiknjo, S. D. et al. Novel Effects of Chromosome Y on Cardiac Regulation, Chromatin Remodeling, and Neonatal Programming in Male Mice. Endocrinology 154, 4746–4756 (2013).

55. Buenrostro, J. D., Wu, B., Chang, H. Y. & Greenleaf, W. J. ATAC-seq: A Method for Assaying Chromatin Accessibility Genome-Wide. in Current Protocols in Molecular Biology (John Wiley & Sons, Inc., 2001).

